# CD8^+^ T cells mediate vaccination-induced lymphatic containment of latent *Mycobacterium tuberculosis* infection following immunosuppression, while B cells are dispensable

**DOI:** 10.1101/2025.01.23.634479

**Authors:** Socorro Miranda-Hernandez, Manoharan Kumar, Alec Henderson, Erin Graham, Xiao Tan, Jim Taylor, Michael Meehan, Zuriel Ceja, Lidia del Pozo-Ramos, Yi Pan, Ellen Tsui, Meg L Donovan, Miguel E Rentería, Mario Alberto Flores-Valdez, Antje Blumenthal, Quan Nguyen, Selvakumar Subbian, Matt A Field, Andreas Kupz

**Author notes:** These authors contributed equally (Manoharan Kumar, Alec Henderson, Erin Graham). Co-senior authors (Selvakumar Subbian, Matt A Field, Andreas Kupz).

## Abstract

It is estimated that two billion people are latently infected with *Mycobacterium tuberculosis* (*Mtb*), the causative agent of tuberculosis (TB). Latent *Mtb* infection (LTBI) can occur in multiple organs, including the lymphatics. The risk of LTBI reactivation increases in immunocompromised conditions, such as coinfection with human immunodeficiency virus (HIV), and during treatment of autoimmune diseases and organ transplantation. The immunological correlates of protection against TB, including against reactivation of LTBI, remain largely elusive. Here, we used a mouse model of latent lymphatic *Mtb* infection to dissect the immunological mechanisms underlying LTBI containment versus reactivation. We show that immunosuppression-mediated reactivation of lymphatic LTBI and the subsequent spread to non-lymphatic organs can be prevented by vaccination with multiple recombinant BCG (rBCG) strains despite the deficiency of CD4^+^ T cells. Using spatial transcriptomics, multi-parameter imaging, network analysis and bioinformatic integration of histopathological images, we reveal that immunosuppression is associated with a distinct repositioning of non-CD4 immune cells at the edge of TB lesions within the infection-draining cervical lymph nodes. While B cells increased in numbers, they are dispensable for the containment of LTBI. Lymphatic *Mtb* infection in different immune cell-deficient mouse strains, antibody-mediated cell depletion and adoptive transfer experiments into highly susceptible mice unequivocally show that vaccination-mediated prevention of LTBI reactivation is critically dependent on CD8^+^ T cells. These findings have profound implications for our understanding of immunity to TB and the management of LTBI.

**Graphical Abstract:** 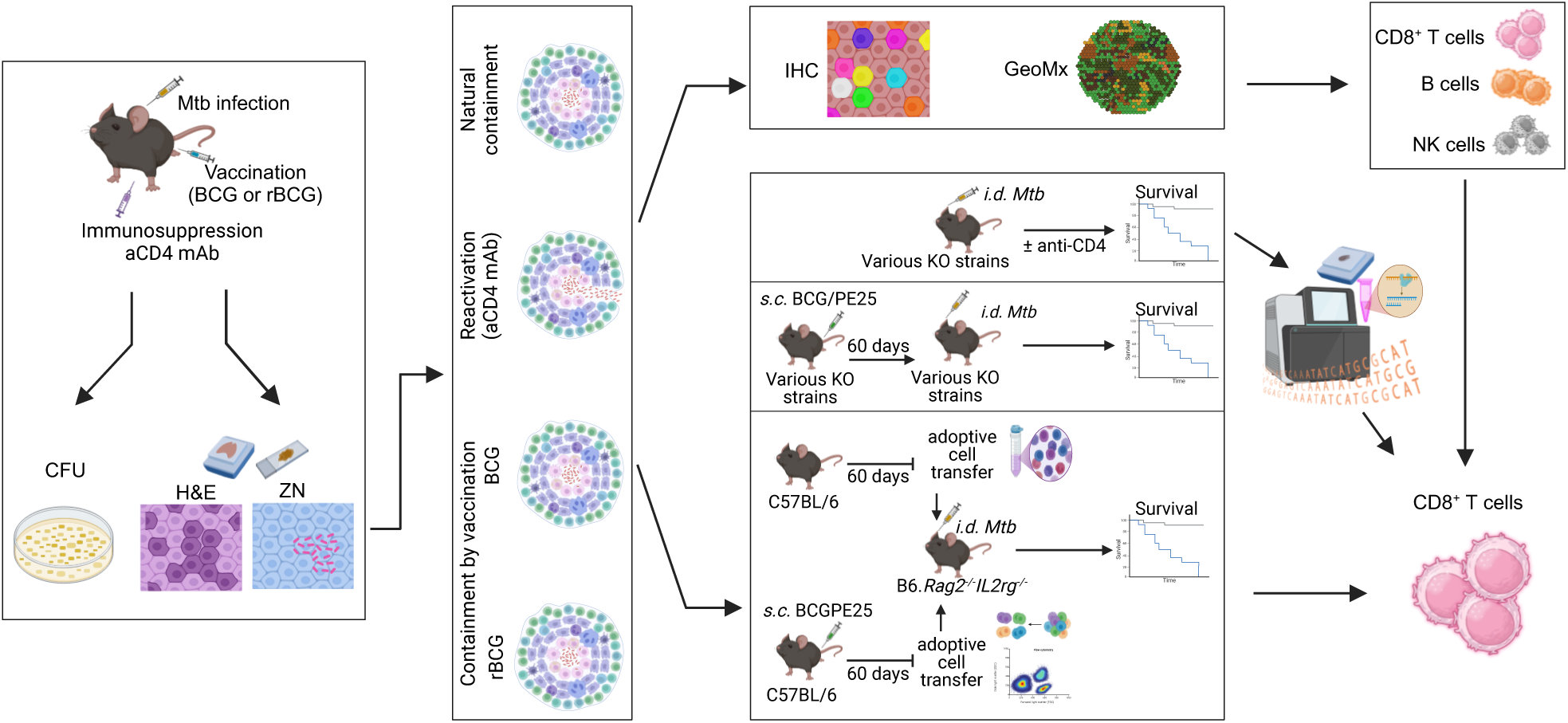

## Introduction

Tuberculosis (TB), caused by *Mycobacterium tuberculosis* (*Mtb*), remains the leading infectious cause of death by a single pathogen globally, with about 1.3 – 1.5 million deaths annually. In addition, it is estimated that about two billion people are latently infected with *Mtb* without showing signs of active disease [1]. Latent *Mtb* infection (LTBI) is often contained within granulomatous lesions that can occur in multiple organs. However, immunosuppressed individuals often fail to control LTBI, and the risk of LTBI reactivation increases with human immunodeficiency virus (HIV) infection, type 2 diabetes, malnutrition, autoimmune diseases, and following organ transplantation [2, 3]. Reactivation of LTBI remains the primary cause of death among individuals co-infected with *Mtb* and HIV. A hallmark of HIV infection and several other immunosuppressive conditions is the impaired function, reduction, or loss of CD4^+^ T cells, crucial immune cells for TB control, the maintenance of the structural integrity of granulomas and a primary target for vaccine-induced protection [4, 5]. However, the specific role of CD4^+^ T cells in latent TB immunity is still debated, especially in immunocompromised individuals [6].

Recognising that LTBI shares features with lymphatic diseases, we previously developed a mouse model to study LTBI reactivation following the loss of CD4^+^ T cells [7]. This murine TB model, which involves ear dermis infection, reflects several immunological aspects of human LTBI. In this model, *Mtb* spreads to the lungs only in mice depleted of CD4^+^ T cells after intradermal (i.d.) infection, while immunocompetent mice contain *Mtb* within the draining lymph nodes (LNs), mimicking latent TB. Historical descriptions of human TB and observations in non-human hosts suggest that TB has lymphatic characteristics, and that pulmonary pathology primarily aids disease transmission [8]. This lymphocentric view of TB is supported by findings that LNs, bone marrow, spleen, thymus, tonsils and adenoids are persistently infected with *Mtb* in humans and non-human primates and serve as a source of LTBI reactivation [9–13]. The ear dermis infection model has also been used to investigate immune responses associated with contained *Mtb* infection more broadly [14, 15].

Importantly, we have previously shown that BCG vaccination can prevent the reactivation of murine LTBI with anti-CD4 monoclonal antibodies (mAb), indicating that vaccine-induced CD4^+^ T cells are not essential to prevent the reactivation of lymphatic murine LTBI [16]. This finding raised the fundamental question of which immune cells control *Mtb* containment within the lymphatics after BCG vaccination in the absence of CD4^+^ T cells. Given the importance of managing and reducing LTBI for the WHO’s End TB Strategy [17], latent murine lymphatic *Mtb* infection offers an opportunity to study LTBI reactivation from a lymphocentric perspective, and provides a valuable platform to explore the immunological requirements for LTBI containment versus reactivation in the context of vaccination. While macrophages and monocytes express CD4 in humans, they do not in mice [18]. This allows the study of CD4^+^ T cell depletion in the presence of macrophages/monocytes. *Mtb* uses alveolar macrophages to disseminate, survive, and replicate; these cells are also critical to initiate the formation of granulomas [19, 20]. Moreover, monocytes are altered in individuals with LTBI, so they may play a role in the reactivation of the disease [21]. In this context, this mouse model selectively mimics the impaired function or loss of CD4^+^ T cells in human LTBI.

Until recently, the dissection of the complex interactions between *Mtb* and the host immune system, which lead to the establishment of LTBI, predominantly relied on histopathological studies and cellular analyses of disrupted tissues or the extraction of DNA and RNA from blood. With the emergence of spatial tissue profiling in conjunction with multiplex imaging and advances in bioinformatics, it is now possible to map and analyse the organisation and activity of thousands, or even millions, of cells, genes, and molecules in the tissue [22]. This approach has been used to map TB granulomas in the lungs, revealing the tissue architecture and cellular interactions in TB. It shows that *Mtb* infected human lung tissue comprises at least four types of non-necrotising structures in addition to necrotising granulomas [23]. Furthermore, spatial transcriptomics has provided unprecedented insights into the pathogenesis of Alzheimer’s disease, Multiple Sclerosis, skin diseases and various types of cancers in different organs [24–27].

Here, we used different experimental strategies to dissect the immunological requirements of lymphatic LTBI containment. One approach utilised the contained *Mtb* reactivation model in conjunction with conventional histopathological and multi-parameter spatial transcriptomics techniques to delineate the LN architecture following lymphatic *Mtb* infection with and without prior BCG vaccination. We investigated whether recombinant BCG (rBCG) strains can prevent the reactivation of lymphatic LTBI and assessed multiple extrapulmonary organs for signs of disseminated TB. In parallel, we performed *Mtb* infections and adoptive cell transfers in various mouse strains lacking distinct immune cell subsets and monitored their ability to resist LTBI reactivation. Our results give detailed high-dimensional insight into the cellular compositions during *Mtb* reactivation and BCG-mediated containment of *Mtb* in the lymphatics. These findings have significant implications for designing preventative and therapeutic strategies to combat LTBI and its reactivation.

## Materials and Methods

### Bacterial cultures

BCG SSI, BCG::ESAT-6-PE25SS, BCGΔBCG1419c::*dosR* (abbreviated to BCG, PE25 and dosR respectively in Figures) [28] and *Mtb* H37Rv were cultured in Middlebrook 7H9 broth enriched with 10% Albumin Dextrose Catalase (ADC) (BD), 0.5% glycerol, and 0.05% Tween 80 (vol/vol). BCG::ESAT-6-PE25SS and *Mtb* H37Rv were grown in the presence of 50 µl/ml kanamycin and 10 µl/ml ampicillin, respectively. The cultures were shaken at 80-90 rpm at 37°C until they reached the log phase with an OD600 of 0.6-0.8. The bacterial cultures were then harvested, washed, aliquoted in 10% glycerol and kept at -80°C until use.

### Mice

Six-to eight-week-old female C57BL/6 mice were obtained from the Animal Resources Centre (ARC) Perth, WA, Australia. B6.*µMT*, B6.*Rag1^-/-^*and B6.*Rag2^-/-^IL2rg^-/-^* mice were bred and maintained in specific pathogen-free conditions at the James Cook University Bioresource Facility of the Australian Institute of Tropical Health and Medicine (AITHM) in Townsville, QLD, Australia. Before performing experiments, the animals were acclimatised to the facility for at least seven days. The mouse rooms were maintained with a 12-hour light/dark cycle, a temperature of 21 ± 2°C, and a humidity of 55% ± 10%. The mice were housed in ventilated cages with bedding consisting of irradiated fine corn cobs, enriched with a cardboard roll, Bed R Nest, cotton, and some pieces of wood for them to chew. Water was treated by reverse osmosis and sterilised by UV light. The irradiated mouse diet was obtained from Specialty Feeds (WA, Australia). Both food and water were provided ad libitum. All experimental procedures were approved by the Animal Welfare and Ethics Committee of James Cook University (approval number A2794). Each mouse was monitored for clinical signs of disease and/or weighed throughout the study.

### Vaccinations and infections

Mice were immunised via subcutaneous (s.c.) injection in the base of the tail with 100µl of phosphate buffer saline (PBS) containing 1×10^6^ CFU of BCG, BCG::ESAT-6-PE25SS or BCGΔBCG1419c::*dosR*. Sixty days after vaccination, mice were immunosuppressed with an intraperitoneal (i.p.) injection of anti-CD4 (aCD4) mAb (clone GK1.5; 200µg/mouse; Walter and Eliza Hall Institute of Medical Research (WEHI) antibody facility) or anti-CD8 (aCD8) mAb (clone 2.43; 200µg/mouse; WEHI antibody facility) and infected intradermally (i.d.) in the ear with 1×10^3^ CFU of *Mtb*. aCD4 and aCD8 antibody injections continued weekly for the duration of the experiment. Naïve, vaccination-only, immunosuppressed-only, and infection-only groups were included as controls. The experimental setup is shown in the Figures. Samples were collected at 30 days, 60 days, or 120 days after *Mtb* infection.

### Disease indicator score

The clinical signs of disease for immunisations, immunosuppression and infections in C57BL/6 mice were scored from 0 to 3 as follows: a) Appearance: 0 = Shiny fur, clear eyes, 1 = Fur appears untended with or without a lesion in the skin, 2 = Scruffy fur, discharge from eyes and nose, eyes sunken and anus dirty or sore, 3 = Very scruffy fur, blood from orifices, abnormal body posture; b) Behaviour/movement: 0 = Normal contact with other mice, active movement, 1 = Little contact with other mice, 2 = Little contact with other mice, avoid movement, 3 = Complete isolation, noises, self-harm, no-movement, lying on side; c) Reaction to stimuli: 0 = adequate, 1 = Slightly delayed, 2 = Significantly delayed or weak, 3 = No reaction; d) Breathing: 0 = Normal; 1 = increased breathing, 2 = increased breathing with sporadic increase in abdominal breathing, 3 = constant abdominal breathing; and e) Weight loss: 0 = none, 1 = Less than 10%, 2 = 10% to 14%, 3 ≥ = 15%. Mice were euthanised when they reached a score of 3 on any criterion.

### Sample collection

Mice were euthanised by cervical dislocation and spleens, cervical LNs and right lung lobes were collected for CFU enumeration. Mesenteric LNs (mLNs), lungs, brains, salivary glands, livers, femurs, cervical LNs, and left lung lobes were fixed in 4% paraformaldehyde for 24 hours. After being fixed, the samples were transferred into 70% ethanol until they were processed for histopathological studies.

### CFU enumeration

Spleens, cervical LNs and lungs were homogenised in Nasco Whirl-Pak bags or gentleMACS tubes containing 1 ml of PBS enriched with 0.05% Tween 80. Tenfold dilutions (N, -1, -2) were plated onto Middlebrook 7H11 agar plates supplemented with 0.02% glycerol, 0.05% Tween 80 and 10% oleic albumin dextrose catalase (OADC). Agar plates were sealed, wrapped in aluminium foil, and incubated aerobically at 37°C, and CFU were counted after four weeks.

### Histopathological studies

Tissue samples were embedded in paraffin and sectioned into 4-micrometre-thick slices. Slices were transferred onto microscope slides, dewaxed, and stained using the following methods:

#### a) Hematoxylin and Eosin (H&E)

mLNs, lungs, brains, salivary glands, livers, femurs, cervical LNs, and left lung lobes were stained with H&E. Briefly, dewaxed tissues were hydrated, stained with hematoxylin, differentiated, blued, and then stained with Eosin. Subsequently, the tissues were dehydrated, cleared, and cover-slipped.

#### b) Ziehl-Neelsen (ZN)

mLNs, lungs, brains, salivary glands, livers, femurs, cervical LNs, and left lung lobes were stained with ZN. In brief, dewaxed tissues were hydrated, stained with Carbol Fuchsin, Ziehl-Neelsen, and differentiated in acid alcohol. Afterwards, tissues were counterstained with methylene blue, dehydrated in ethyl alcohol, cleared, and cover-slipped.

#### c) 9-colour Opal Immunohistochemistry (IHC)

Cervical LNs and lungs were stained with a 9-colour Opal IHC Multiplex panel (Akoya) by the Advanced Histotechnology Facility of the WEHI, Melbourne, Victoria, Australia. The 9-colour opal IHC panel included the following antibodies: *Mtb* (Cat. No. ab905, clone pAb, Abcam), CD3 (Cat. No. A045229-2, clone pAb, Dako), CD4 (Cat. No. 14-9766-82, clone 4SM95, Invitrogen), CD8a (Cat. No. HS361-008, clone Rb321E9, Synaptic systems), B220 (antibody from WEHI), CD161c/NK1.1 (Cat. No. bs-4682R, clone pAb, biossusa), CD11c (Cat. No. 97585, clone D1V9Y, Cell Signalling Technology), CD11b (Cat. No. ab133357, clone EPR1344, Abcam), and Spectral DAPI (Cat. No. FP1490, Akoya Bioscience). The 9-colour Opal IHC Multiplex samples were analysed using QuPath (versions 3 to 5).

#### d) Whole Transcriptome Analysis (WTA)

Cervical LNs were stained using the NanoString GeoMx Mouse Whole Transcriptome Assay (version 3.0.0.182) by NanoString Technologies, Inc. (Seattle, Washington, USA). Thirty regions of interest (ROIs) were selected based on areas with different histopathological changes seen in H&E. These ROIs included three pathological areas: I) Normal in Naïve = no pathological changes in mice that did not receive treatment, and Normal in disease = no pathological changes were seen in tissues from *Mtb*-only infected mice, immunosuppressed and *Mtb*-infected, and vaccinated-immunosuppressed and *Mtb*-infected; II) Lesion = tissue resembling granuloma; and III) Edge of the Lesion = border of the tissue resembling granuloma. Cellular deconvolution for estimating the abundance of cell types within each ROI was performed using the GeoMx DSP Analysis Suite. The analysis utilised the DSPPlugSpatialDecon and the scripting file SpatialDecon_plugin.R alongside the cell profile metrics ImmuneAtlas_cellFamily_ImmGen.

### Digital image analysis

QuPath versions 3.1-5.1 were used to analyse H&E, ZN and Multiplex IHC slides. Each LN and lung lobe were selected for the three different staining techniques. The areas affected by *Mtb* were identified, and their respective percentages were calculated for H&E. In the ZN staining, *Mtb* were identified, and the proportion of the area occupied by the bacteria was calculated. In the multiplex IHC, cell nuclei and segmentation were identified using DAPI. Subsequently, two methodologies were applied. Briefly, each cell was determined by a threshold and classified according to the marker expressed by a threshold, and subsequently, multiple markers were used to train a machine-learning classification of the cell types for each cell. Once each cell was classified according to its phenotype, cells were enumerated, and distances among cells were analysed [29].

### Cell isolation and adoptive transfer

Spleen and LNs were collected aseptically from naïve and vaccinated C57BL/6 donor mice. The tissues were dissociated using a sterile three cc syringe plunger and passed through a 70μm cell strainer. Red cells were lysed using red cell lysis buffer.

To deplete CD3^+^ and/or CD19^+^ cells, LNs were disrupted as described above. The cell separation was carried out using the CD3e MicroBead kit and CD19 MicroBeads (Miltenyi Biotec). Briefly, single-cell suspensions were labelled with CD3-or CD19-conjugated immunomagnetic beads and then depleted using a MACS system (Miltenyi Biotec). Further enrichment was achieved by passing the negative fraction through a second MACS column. Flow cytometry analysis showed that approximately 90-98% of CD3^+^ and/or CD19^+^ cells were effectively removed.

For adoptive transfer experiments [30], B6.*Rag2^-/-^IL2rg^-/-^* recipient mice received 5.7 ×10^5^ to 7 ×10^6^ cells via intravenous injection into the tail vein. The recipient mice were infected with *Mtb* intradermally 1-7 days after cell transfer.

### RNA sequencing from fixed cervical lymph nodes

RNA extraction, library preparation, sequencing and initial data processing were performed by Lexogen NGS Services, Lexogen GmbH, Austria. In brief, RNA was isolated using PureLink™ FFPE RNA Isolation Kit (ThermoFisher). Samples were characterized by UV-Vis spectrophotometry (Nanodrop2000c, Thermo Fisher) and RNA integrity was assessed on a Fragment Analyzer System using the DNF-471 RNA Kit (15 nt) (Agilent). To remove genomic DNA contaminants, the samples were treated with DNase I. The libraries were constructed using Lexogen’s QuantSeq 3’ mRNA-Seq Library Prep Kit FWD. Prepared libraries were quality controlled on a Fragment Analyzer System using the DNF-474 using the HS-DNA kit (1-6000 bp) (Agilent). cDNA libraries were sequenced with an Illumina NextSeq 2000 platform using 100 nt single end read length, with 5 million reads on average per sample.

Sequencing quality control of the raw reads was assessed using FastqQC software and adapter sequences were removed with cutadapt [31] Alignment to the *Mus musculus* reference genome (GRCm38, Ensembl release 102) and read counting were performed using STAR [32] and featureCounts [33]. The DESeq2 R package [34] was used to perform differential gene expression analysis.

### Data analysis & statistics

Data analysis was performed using different software; GraphPad Prism versions 9.3.1 to 10.1.2; R and RStudio version 4.3.1. R packages included NanoStringNCTools, GeomxTools, GeoMxWorkflows and GeoMx-NGS gene panel version Mm_R_NGS_WTA_v1.0. For image analysis and representative images, QuPath versions v0.3.1 to v0.5.0, Image J version 1.53 and GIMP versions 2.10.30 to 2.10.36 were used. The significance of differentially expressed genes (DEGs) in the volcano plots was determined using a false discovery cut-off rate (FDR) < 0.05. The Venn diagrams were plotted using significant DEGs for each comparison with the VennDiagram package in R and QIAGEN CLC Genomic Workbench version 24.0.1. The pathway analyses were done using the QIAGEN Ingenuity pathway analysis (IPA) 2023-2024 releases. Results are presented as data means ± SEM. Statistical comparisons were performed using One-way ANOVA with Tukey’s multiple comparison test, Kruskal-Wallis test with Dunn’s multiple comparison test, or Long-rank (Mantel-Cox) test. Data were considered significant when \**p* < 0.05, ** *p* < 0.01, *** *p* < 0.001, **** *p* < 0.0001.

## Results

### BCG and rBCG vaccination prevent *Mtb* dissemination following reactivation of lymphatic LTBI

rBCG strains have attracted considerable attention as potential substitutes for BCG [35, 36]. We and others have developed candidate TB vaccines by integrating immunogenic proteins from the Region of Difference 1 (RD1) locus of *Mtb* into BCG. The first-generation rBCG strains containing RD1 from *Mtb* were considered inappropriate to progress to human trials due to prolonged persistence and increased virulence in immunocompromised hosts [37]. However, next-generation rBCG candidates, such as *Mtb*::ESX1^Mmar^ and BCG::ESAT-6-PE25SS [38, 39] aim to maintain the enhanced immunogenicity while preventing the detrimental aspects of *Mtb*-derived RD1. To enhance the likelihood of potential translation to clinical trials, it is important to evaluate rBCG strains in different pre-clinical models of TB.

To determine if BCG::ESAT-6-PE25SS can also prevent the reactivation of lymphatic LTBI and the spread of *Mtb* to non-lymphatic organs, six-to eight-week-old C57BL/6 mice were grouped and treated as described in the Materials and Methods section and illustrated in **Fig. 1A**. Mice did not show signs of disease (**Fig. 1B**), and although all the mice gained weight throughout the experiment (**Fig. 1C-E**), a comparison between vaccinated and naïve mice revealed a slight delay in weight gain, particularly from week 20 to week 26 (**Fig. 1C**). BCG-vaccinated mice infected with *Mtb* experienced delayed weight gain during weeks 24 and 25, compared to mice only infected with *Mtb* (**Fig. 1D**). Both BCG-vaccinated, anti-CD4 mAb-treated and *Mtb*-infected mice, as well as those treated with anti-CD4 mAb and infected with *Mtb*, experienced a delay in weight gain for a week (week 25 and week 26 respectively) compared to mice only infected with *Mtb* (**Fig. 1E**).

**Figure 1.**
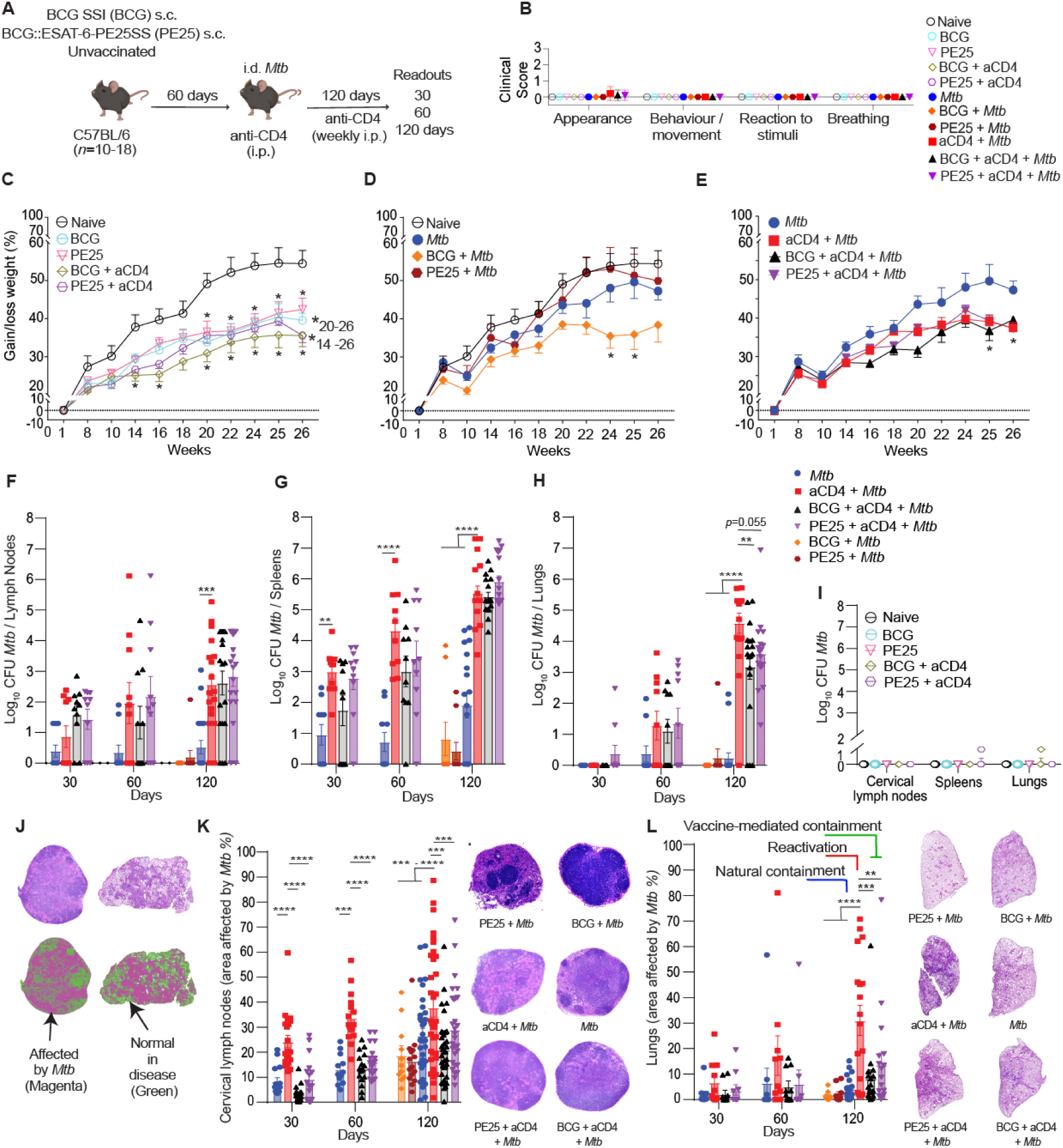
rBCG vaccination prevents pulmonary TB following the reactivation of LTBI via immunosuppression. **(A)** Experimental setup. C57BL/6 mice were vaccinated s.c. with 1×10^6^ CFU of either BCG, BCG::ESAT-6-PE25SS (PE25) or left unvaccinated. Sixty days after vaccination, mice were infected i.d. with 1×10^3^ CFU of *Mtb* H37Rv. Some mice received anti-CD4 mAb (referred to in figure as aCD4) weekly i.p. for 120 days. Groups without *Mtb* infection or anti-CD4 mAb treatment were included as controls. **(B)** Clinical scoring of disease, and **(C-E)** percentage of body weight gain/loss over time. **(F-H)** CFU of *Mtb* in cervical LNs **(F)**, spleens **(G),** and lungs **(H)** at 30, 60 and 120 days post *Mtb* infection. **(I)** CFU of untreated (naïve), vaccination-only and uninfected and anti-CD4 mAb-treated controls in cervical LNs, spleens and lungs at 120 days. **(J)** Representative H&E sections of cervical LNs (left) and lung left lobes (right) showing areas affected by *Mtb*. Normal tissue in disease (green) and areas affected by *Mtb* (magenta). **(K)** Percent of tissue affected by *Mtb* in cervical LNs over time and representative H&E sections from each treatment group. **(L)** Percent of tissue affected by *Mtb* in the left lung lobes over time indicating natural containment (blue), reactivation (red), and vaccine-mediated containment (green). Representative H&E sections of left lung lobes from each treatment group. Results are presented as data means ± SEM **(B-E)**. Individual data points ± SEM (**B, F-I, K, L**) and representative images **(J-L)** from 3 pooled independent experiments (*n* = 10-18 mice per group). Statistical analyses: One-way ANOVA followed by Tukey’s multiple comparison test per time-point or Kruskal-Wallis test with Dunn’s multiple comparison test; asterisks indicate significant differences: * *p* < 0.05, ** *p* < 0.01, *** *p* < 0.001, **** *p* < 0.0001. See also Figure S1.

Mice that only received *Mtb* infection contained *Mtb* in the lymphatics (cervical LNs and spleen) with negligible *Mtb* dissemination to the lungs, particularly at later time points (**Fig. 1F-H**). As previously shown [7, 16], anti-CD4 mAb treatment led to a progressive increase in bacterial burden in LNs, spleens and lungs, with the most pronounced level of reactivation seen at 120 days after *Mtb* infection (**Fig. 1F-H**). While vaccination with BCG and BCG::ESAT-6-PE25SS only marginally reduced the level of *Mtb* bacilli in the lymphatic organs relative to anti-CD4 mAb-treated mice, it significantly reduced and, in some cases, even prevented the spread of bacteria to the lungs (**Fig. 1F-H**). All control groups that did not receive *Mtb* showed no detectable *Mtb* bacteria in any of the organs at the 120-day time-point (**Fig. 1I**). Strikingly, when cervical LNs and lungs were assessed for histopathological changes using the percent of tissue area covered by positive signal, implementing the automated positive pixel function in QuPath (**Fig. 1J-L**), it became obvious that BCG and BCG::ESAT-6-PE25SS vaccination had a dramatic effect on tissue morphology and significantly reduced measurable pathology from day 30 onwards in the LNs (**Fig. 1K**), and led to a significant reduction of lung pathology at 120 days after infection (**Fig. 1L**). Representative images show the area affected at 120 days after *Mtb* infection in cervical LNs and lungs (**Fig. 1K, L**). Using a third rBCG strain, BCGΔBCG1419c::*dosR* (abbreviated to dosR in **Fig. S1A-C**), we confirmed the significant reduction of lung histopathology at day 120 (**Fig. S1A-C**). Collectively, those results indicate that vaccination with BCG or rBCG strains significantly reduces the spread of *Mtb* from the lymphatic system to the lung and prevents the induction of lung pathology in this model of LTBI reactivation. Hence, for subsequent analyses we will refer to three distinct scenarios: i) natural containment (*Mtb* infection only); reactivation (*Mtb* infection plus anti-CD4 mAb treatment); and vaccine-mediated containment (vaccination followed by *Mtb* infection and anti-CD4 mAb treatment) (illustrated in **Fig. 1L**).

### LTBI reactivation occurs across multiple lymphatic and non-lymphatic organs

Given that TB can occur in multiple organs [40], we investigated whether reactivation of LTBI in our model impacts additional lymphatic and non-lymphatic organs and whether vaccination can prevent this. To this end, histopathological analyses and automated scoring were conducted on mLNs, femurs, livers, brains, and salivary glands. Like in the cervical LNs (**Fig. 1K**), anti-CD4 mAb treatment led to significantly enhanced histopathology in mLNs from 60 days after *Mtb* infection. Vaccination with BCG and BCG::ESAT-6-PE25SS markedly and significantly reduced the level of mLN damage (**Fig. 2A**). Interestingly, the level of histopathological damage observed in the liver was similar across all groups and all time points regardless of vaccination or anti-CD4 mAb treatment but above baseline naïve levels for all groups (**Fig. 2B**). When femurs were assessed for histopathology, we could not observe any changes to the bone structure at any of the time-points measured; however, the affected area of bone marrow increased with anti-CD4 mAb treatment from day 60, and it reached significant levels of damage by 120 days compared to the *Mtb*-only group (**Fig. 2C**). There was no significant reduction of bone marrow damage in the vaccinated groups that also received anti-CD4 mAb treatment. No pathological changes were observed in either the brain or the salivary glands in any of the treatment groups at 120 days after infection (**Fig. 2D**). In line with these histopathological results, we also found significantly increased tissue area occupied by *Mtb* in the cervical LNs, the lungs, the mLNs, the livers, and the bone marrow of anti-CD4 mAb-treated animals using Ziehl-Neelsen staining (**Fig. 2E**). Both BCG strains significantly reduced this area in all those tissues to the level of *Mtb*-infected mice that did not receive anti-CD4 mAb treatment. Collectively, these results show that anti-CD4 mAb-mediated reactivation of LTBI occurs across multiple lymphatic organs, the lung and the liver, but not the brain or the salivary glands, and that vaccination can prevent this spread.

**Figure 2.**
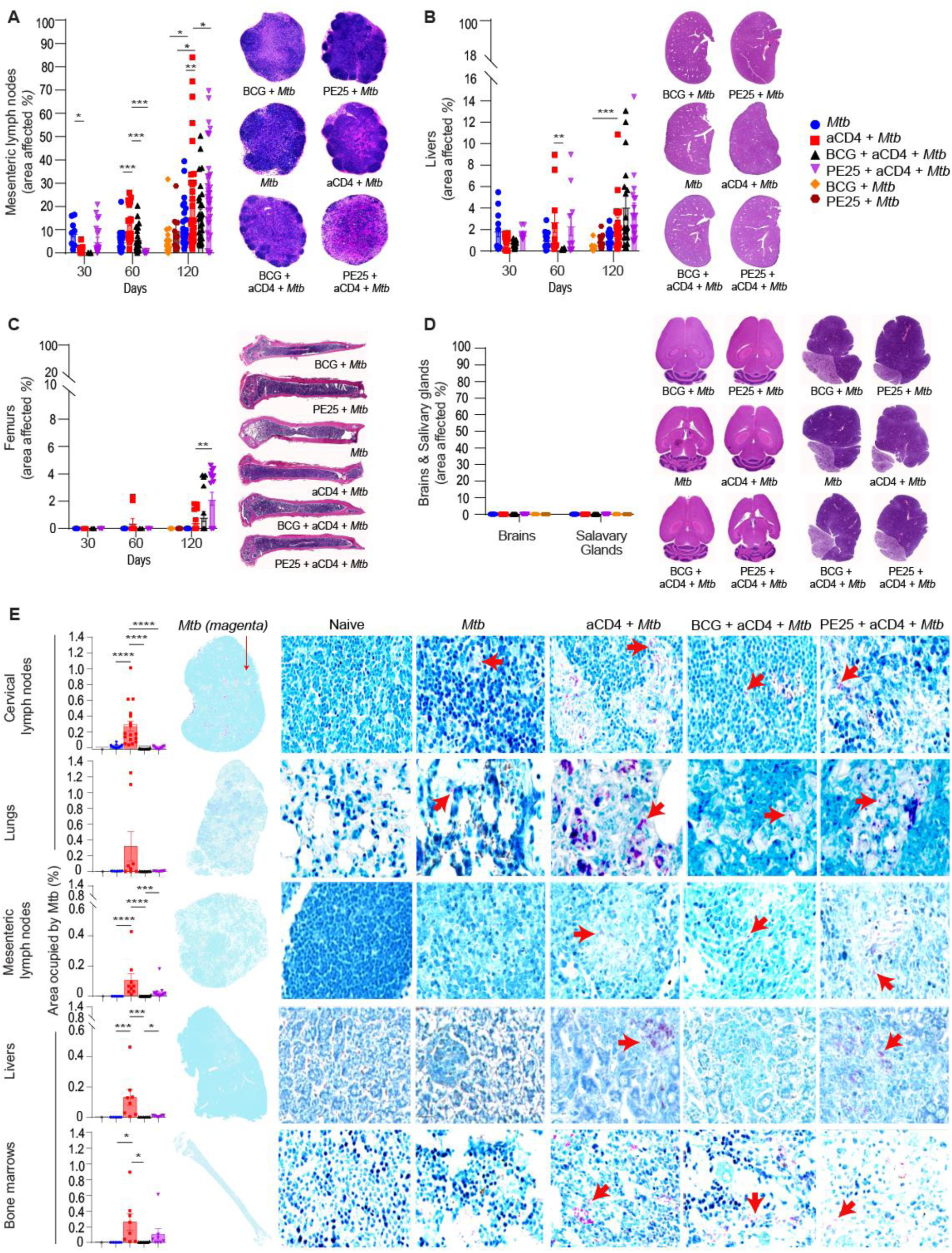
LTBI reactivation occurs across multiple lymphatic and non-lymphatic organs. (**A-D)** Percent cellular infiltration in mesenteric LNs **(A)**, livers **(B)**, femurs **(C)**, salivary glands and brains **(D)** over time, including a representative H&E-stained section of each tissue from different treatment groups. (**E)** Percentage of the area occupied by *Mtb* in cervical LNs, lungs, mLNs, livers, and bone marrows with representative sections stained with Ziehl-Neelsen (ZN). Bacteria are shown in magenta and indicated with red arrows. Results are presented as individual data points ± SEM **(A-E)** and representative images from 3 pooled independent experiments (*n* = 8-18 mice per group). Statistical analyses: One-way ANOVA followed by Tukey’s multiple comparison test per time-point; asterisks indicate significant differences: * *p* < 0.05, ** *p* < 0.01, *** *p* < 0.001, **** *p* < 0.0001.

### Containment of lymphatic LTBI is associated with cellular changes in the infected LNs and repositioning of immune cells

To determine the immunological mechanisms that may be associated with natural containment (*Mtb*-only) vs. vaccination-mediated containment (BCG or rBCG with anti-CD4 mAb) vs. anti-CD4 mAb treatment-mediated reactivation of lymphatic LTBI, we first performed multiplex immunohistochemistry (IHC) using eight fluorescent antibodies against CD3, CD4, CD8, B220, CD11b, CD11c, CD161c, *Mtb* and DAPI for the nuclei (**Fig. 3A-C**). Enumeration of single positive cell types across the entire cervical LNs confirmed an increase in *Mtb*-infected cells in the anti-CD4 mAb-treated group relative to the *Mtb*-only group and groups that received anti-CD4 mAb treatment but had been vaccinated prior with BCG or BCG::ESAT-6-PE25SS (**Fig. 3D**; **Fig. S2A).** Multiplex IHC also confirmed the efficient depletion of CD4^+^ cells in all groups that had received anti-CD4 mAb (**Fig. 3D**). Similar observations occurred when single positive cells of each marker were enumerated across the whole lung lobes (**Fig. 3E**; **Fig. S2B**).

**Figure 3.**
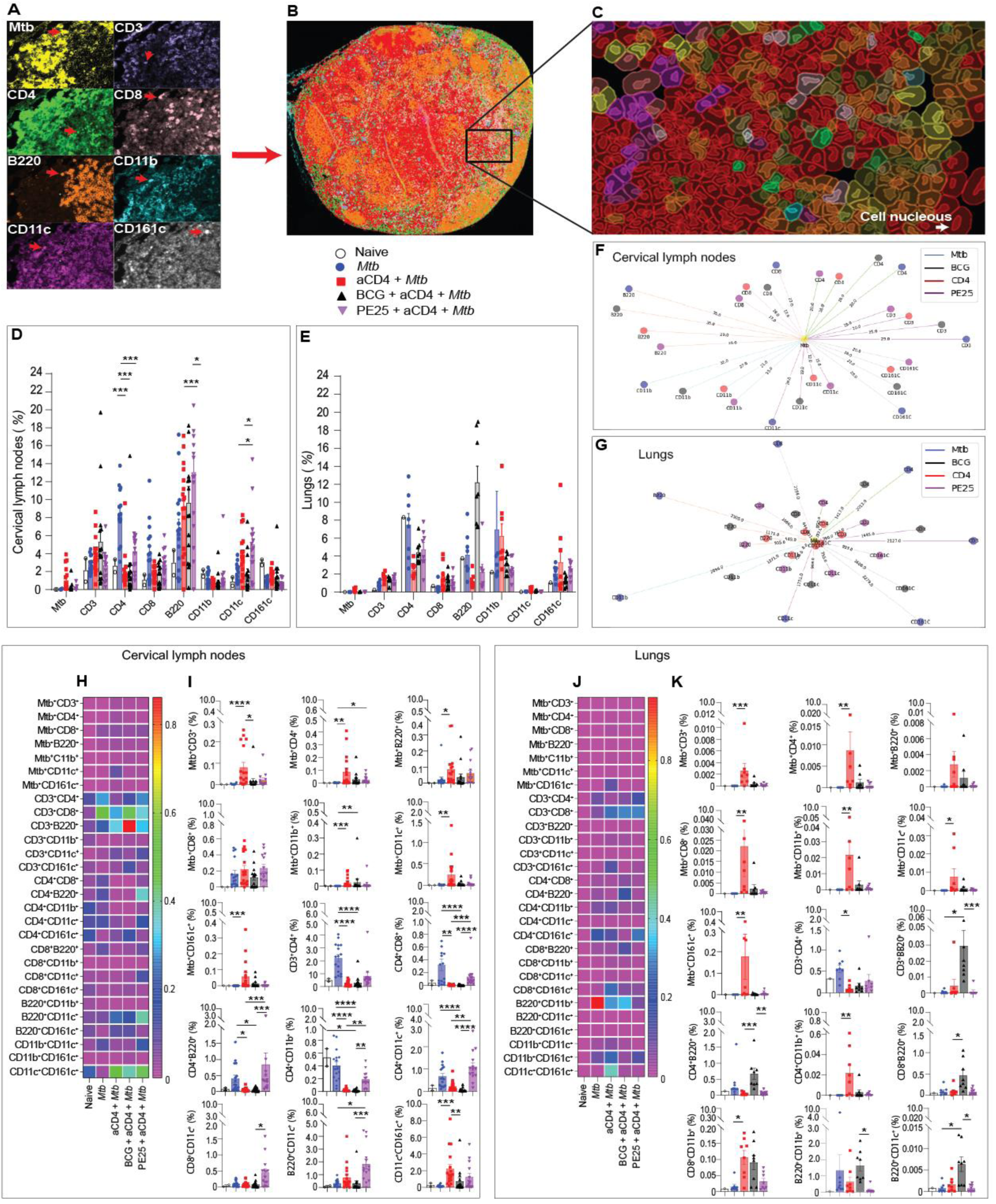
BCG vaccination increases B cell numbers and alters cellular positioning around *Mtb-* infected cells. **(A)** Representative images of Opal multiplex immunohistochemistry obtained from LN samples. **(B)** Representative LN image after classifying each cell type with its respective marker. **(C)** Enlarged area delineating the cells and nucleus of each cell, coloured according to the markers expressed within them *Mtb* (yellow), CD3 (purple), CD4 (green), CD8 (pink), B220 (orange), CD11b (cyan), CD11c (magenta) and CD161c (white). (**D**, **E**) Percentage of single positive cells across entire cervical LNs **(D)** and left lung lobes **(E)**. (**F**, **G**) The median distance in micrometres of CD3, CD4, CD8, B220, CD11b, CD11c or CD161c positive cells to *Mtb* infected cells in LNs **(F)** and left lung lobes **(G).** Heatmaps showing double positive cell subtypes of all combinations seen across entire LNs **(H)** and left lung lobes **(J)**. Percentage of cells expressing two markers in cervical LNs **(I)** and lungs **(K)** with significant differences across groups. Results are presented as individual data points ± SEM **(D, E, I, K)** and representative images **(A-C)** from 2-14 LNs derived from 2 independent experiments at 120 days after *Mtb* infection (n = 8 mice per experimental group; 2 naïve controls). Statistical analyses: One-way ANOVA followed by Tukey’s multiple comparison test per cell type; significant differences are indicated by asterisks: * *p* < 0.05, ** *p* < 0.01, *** *p* < 0.001, **** *p* < 0.0001. See also Figures S2-S4.

We subsequently extended the image analysis to determine cells positive for two of the eight cellular markers used in the Multiplex IHC panel (**Fig. 3H-K**). In the cervical LNs and the lungs, *Mtb*-infected cells were increased in the anti-CD4 mAb-treated group across all the double-positive populations except *Mtb*^+^CD8^+^ in LN and *Mtb*^+^B220^+^ in the lungs (**Fig. 3H-K**). In line with the above results, it was confirmed that CD4^+^ T cells were depleted, as indicated by the reduction of cells expressing CD3^+^CD4^+^ in the anti-CD4 mAb-treated groups in both organs compared to the *Mtb*-only group (**Fig. 3H-K**).

In the LNs, the number of CD4^+^B220^+^, CD4^+^CD11b^+^ and CD4^+^CD11c^+^ double-positive cells also decreased in anti-CD4 mAb treated mice, except those vaccinated with rBCG. In the lungs, CD4^+^B220^+^ and CD4^+^CD11c^+^ cells increased with BCG vaccination and CD4^+^CD11b^+^ cells with anti-CD4 mAb reactivation. Natural containment in the LNs and anti-CD4 mAb-mediated reactivation in the lungs led to a significant increase of double-positive CD4^+^CD11b^+^ cells but no such increase was seen in the groups treated with BCG or rBCG (**Fig. 3I, K**). In contrast, CD3^+^B220^+^, CD4^+^B220^+^, CD8^+^B220^+^ and B220^+^CD11c^+^ cells showed a significant rise in the BCG-vaccinated group in the lungs (**Fig. 3K**). Natural containment and rBCG vaccination in the LNs and anti-CD4 mAb-mediated reactivation in the lungs also led to a significant rise in double-positive CD3^+^CD8^+^ cells (**Fig. 3I, H**). Additionally, CD161c^+^CD11c^+^ cells increased after anti-CD4 mAb treatment and BCG vaccination in the LNs but did not show a significant increase in the lungs (**Fig. 3I, K**). Collectively, these results correlated the increased *Mtb* burden observed in the anti-CD4 mAb-treated group with CFU plating (**Fig. 1F, H**), H&E (**Fig. 1K, M**) and ZN histopathology (**Fig. 2E**). The findings suggest that control of lymphatic LTBI through vaccination may be linked to either the transition or influx of B cells into the lungs, or to changes in the functionality of CD8^+^ T cells in the LNs.

To measure if *Mtb* containment in the LNs and lungs is also associated with an altered immune cell localisation around *Mtb-*infected cells, we performed proximity measurements whereby the distance of all cells was calculated relative to *Mtb*-infected cells. By analysing the distance of over 50 million cells expressing one to eight different markers in relation to *Mtb*-infected cells, it was found that cells expressing multiple markers were fewer and were located closer to *Mtb* in both LNs and lungs (**Fig. S3)**. In LNs, a noticeable shift of all single-positive cells towards *Mtb* in the anti-CD4 mAb-treated reactivation group was observed (**Fig. 3F**; **Fig. S4A**). In contrast, LTBI containment in the *Mtb*-only group and the BCG-vaccinated group was associated with similarly larger cellular distances from *Mtb*, while the anti-CD4 mAb-treated group clustered together with the BCG::ESAT-6-PE25SS-vaccinated group (**Fig. 3F**; **Fig. S4A**). Even though CD8^+^ cells did not increase in the LNs following anti-CD4 mAb treatment (**Fig. 3D**) they showed a consistent pattern of the shortest distance from *Mtb* in *Mtb*-only and BCG-vaccinated groups. In contrast, CD11c^+^ cells were closest to *Mtb* in the reactivation and rBCG-vaccinated groups (**Fig. 3F**; **Fig. S4A**). Surprisingly, B220^+^ cells showed the largest distance to *Mtb* in all groups followed by CD11b^+^ in *Mtb*-only, anti-CD4 mAb reactivation and rBCG-vaccinated groups. CD4^+^ cells were closer to *Mtb* in the BCG-vaccinated group. CD3^+^ and CD161c^+^-expressing cell types maintained an intermediate distance to *Mtb* (**Fig. 3F, G**; **Fig. S4A, B**). A similar pattern was observed when the distance of double-positive cell types relative to the *Mtb*-infected cells was measured (**Fig. 3G**; **Fig. S4A).**

In the lungs, like the LNs, there was a noticeable shift towards *Mtb* in all single-positive cells within the anti-CD4 mAb reactivation group (**Fig. 3G and Fig. S4B).** Although the localisation of each type of cell varied according to the organ, CD8^+^ cells were again closer to *Mtb,* and B220^+^ were farthest away from *Mtb*. CD161c^+^ cells were closer only in the anti-CD4 mAb reactivation group. The distances of cells to *Mtb*, like cell numbers, seem to be organ specific. It appears that natural-and BCG-mediated containment of lymphatic LTBI is linked to a repositioning of CD8^+^ cells around *Mtb* in the infected LNs and lungs, while CD161c^+^ localise around *Mtb* during LTBI reactivation. Collectively, these results suggest a potential protective role for CD8^+^ cells during LTBI containment, a role of CD161c^+^ during reactivation and a potentially permissive role for B220^+^ cells.

### Spatial transcriptomics reveals distinct pathways and regulators of LTBI containment and reactivation

To assess whether LTBI reactivation and vaccine-mediated containment are associated with distinct transcriptomics signatures within the cervical LNs, we performed NanoString GeoMx-based spatial transcriptomics across 30 ROIs that were selected within normal tissue (in naïve and in disease), the lesion or the edge of a lesion from cervical LNs across all treatment groups (**Fig. 4A**). A nonlinear dimensional reduction analysis, namely Uniform Manifold Approximation and Projection (UMAP) showed some clustering of ROIs based on tissue type and treatment group (**Fig. 4B**). When DEGs were ranked and visualised using a heatmap, as expected we observed relatively minor changes in ROIs derived from normal tissue, although the ‘normal tissue in disease’ differed to some degree from normal LN tissue derived from naïve mice (**Fig. 4C**). Across all 30 ROIs there was no overlap of DEGs between *Mtb* infection and anti-CD4 mAb-mediated reactivation relative to naïve mice (**Fig. 4D**). When DEGs were visualised as volcano plots passing the statistical threshold of p<0.05 (orange) and a false discovery rate of <0.05 (blue), distinct DEGs associated with naïve tissue, *Mtb*-based containment, anti-CD4 mAb-mediated reactivation and vaccine-mediated containment were identified (**Fig. 4E-H**). Genes with a *p*<0.05 from the volcano plots were used to analyse the biological pathways and the genes regulating these pathways (**Fig. 4I-N**). Pathways identified from our data by Ingenuity Pathway Analysis (IPA) revealed neutrophil degranulation as the top-upregulated pathway for anti-CD4 mAb reactivation and BCG vaccination. Pathway analysis also revealed that both natural and vaccination induced LTBI containment are associated with keratinisation. However, the pathways ranked just below neutrophil degranulation differentiated anti-CD4 mAb reactivation from BCG vaccination. In particular, class I MHC-mediated antigen processing and presentation and NF-kβ signalling were associated with anti-CD4 mAb-mediated reactivation but not with BCG vaccination (**Fig. 4I**; **Fig. S5A).** The upregulated pathways associated with natural LTBI containment were G-protein coupled receptor signalling and S100 family signalling pathways, respectively (**Fig. 4I**; **Figs. S5A; S7A)**. The upregulated pathways associated with rBCG vaccination were EPH-Ephrin signalling, and G-protein coupled receptor signalling (**Fig. 4I-K**; **Figs. S5A; S7B)**. The pathway regulators were distinct in each treatment group (**Fig. 4J**).

**Figure 4.**
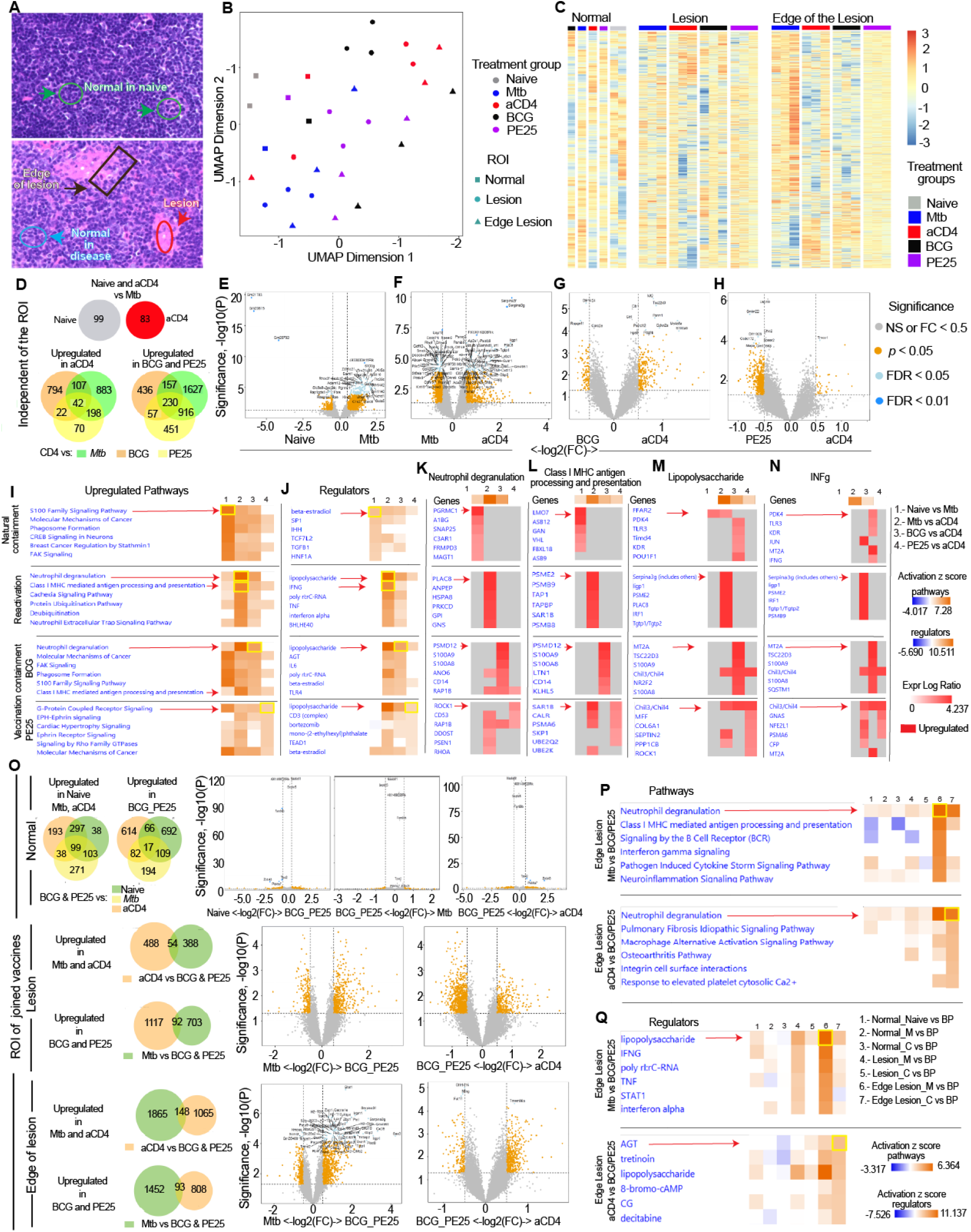
Transcriptomic changes due to reactivation or containment of LTBI are associated with the edge of a lesion. NanoString GeoMx spatial transcriptomics results were derived from 30 regions of interest (ROIs). **(A)** Representative H&E-stained sections from cervical LNs showing the different ROI types used in the study. Normal in naïve (green) and during disease (blue), lesion (red), and edge of the lesion (black). **(B)** UMAP analyses show clustering of ROIs based on tissue type and treatment group. **(C)** Heatmap showing differentially expressed genes (DEGs) across the different ROIs and treatment groups. (**D)** Venn diagrams showing unique and shared upregulated genes in the different treated groups independent of the ROIs. **(E-H)** Volcano plots showing DEGs passing statistical thresholds (orange) and false discovery (blue) across all 30 ROIs and groups. Naïve was compared to *Mtb*-only (**E**), *Mtb*-only was compared to anti-CD4 mAb-mediated reactivation (**F**), anti-CD4 mAb-mediated reactivation was compared to BCG vaccination-mediated containment (**G**) and anti-CD4 mAb-mediated reactivation was compared to BCG::ESAT-6-PE25SS vaccination-mediated containment (**H**). (I-N) IPA predicted upregulated pathways, the six top IPA upregulated pathways in natural containment, anti-CD4 mAb-mediated reactivation and BCG and rBCG-vaccination-mediated containment (**I**), the six top regulators of the highest pathways in each condition **(J)**, the highest expressed genes in neutrophil degranulation **(K)**, the highest expressed genes in Class I MHC antigen processing and presentation **(L)**, top genes upregulated in lipopolysaccharide-driven responses **(M)**, and top expressed genes in IFNγ-driven pathways **(N). (O-Q)** Transcriptomic changes associated with specific ROI types: normal, lesion and edge of the lesion. **(O)** Venn diagrams and volcano plots showing the comparisons of the ROIs in normal, lesion and edge of the lesion across the treatment groups relative to the combined vaccines BCG and BCG::ESAT6-PE25SS referred to in the figure as BP; anti-CD4 referred to as C; *Mtb*-only referred to as M. IPA predicted pathways **(P)** and their regulators **(Q)** at the edge of the lesion. Yellow boxes and red arrows indicate top-regulated pathways and regulators. Results are presented as heatmaps **(C, I-N, P, Q)**, Venn diagrams **(D, O)**, volcano plots **(E-H, O)** and representative images **(A)** from 2-14 LNs derived from 2 independent experiments at 120 days after *Mtb* infection (n = 8 mice per experimental group; 2 naïve controls). Statistical analyses as described in Materials and Methods. The significance of differentially expressed genes in the volcano plots was performed using a false discovery cut-off rate (FDR) < 0.05. The Venn diagrams were plotted using significant DEGs for each comparison with the Venn Diagram package in R. See also Figures S5-S8.

Although lipopolysaccharides were the top predicted regulators of neutrophil degranulation in both the anti-CD4 mAb reactivation and BCG-vaccination groups, IFNγ, class I MHC antigen presentation and processing as well as angiotensinogen (AGT), respectively differentiated anti-CD4 mAb reactivation from the BCG vaccination group (**Fig. 4K-N**; **Fig. S6A)**. Additionally, immunoglobulin in anti-CD4 mAb reactivation and lipopolysaccharides in BCG vaccination were identified as regulators, highlighting distinct mechanisms underlying reactivation and BCG-mediated containment **(Fig. S5B)**.

Collectively, many pathways that were upregulated in the anti-CD4 mAb reactivation group were downregulated or unchanged in the *Mtb*-only and in both vaccination groups, further differentiating BCG-mediated containment of LTBI on a transcript level. Strikingly, in line with our observations that B cells and CD8 cell numbers and distribution were altered following anti-CD4-mediated reactivation of LTBI (**Fig. 3**), immunoglobulins and MHC-I were also identified in the pathway analysis differentiating *Mtb* containment from anti-CD4 mAb-mediated reactivation on a transcriptional level (**Figs. S5B; S6B**).

Given that both BCG and BCG::ESAT-6-PE25SS showed similar patterns of transcriptional changes, we also combined both groups into one vaccination group (BCG_PE25) and compared ROIs across tissue types. In this scenario, as expected, fewer DEGs passed the false discovery rate of <0.05 (**Fig. 4O**). Still, many significant DEGs differentiated anti-CD4 mAb-mediated reactivation from vaccine-mediated containment in the lesion and at the edge of a lesion (**Fig. 4O-Q**; **Fig. S8A-B**). Notably, most of the DEGs passing the false discovery rate were identified at the edge of a lesion rather than within the lesion (**Fig. 4O**). Interestingly, neutrophil degranulation, class I MHC-mediated antigen processing and presentation, lipopolysaccharides, immunoglobulins and AGT genes were all associated with the edge of the lesion (**Fig. 4P-Q**; **Fig. S8A-B**). Collectively, these results revealed that transcriptomic changes associated with reactivation and containment of lymphatic LTBI are unique and predominantly localised at the edge of a lymphatic lesion.

### Cellular deconvolution confirms distinct cellular patterns during containment and reactivation

We used the spatial transcriptomic data and NanoStringNCTools to perform transcript-based cellular deconvolution across all ROIs (**Fig. 5A**). In accordance with our key findings from the Multiplex IHC analysis (**Fig. 3**), we observed significant changes in the proportions of several cell types: T cells (*cd3*), B cells (*cd19*), CD8 cells (*cd8*), CD4 (*cd4*), dendritic cells (*Itgax*), and NK cells (*Klrblc*). These changes distinguished LTBI containment from anti-CD4 mAb-mediated reactivation (**Fig. 5A-H**). Particularly the scaled abundance of identified cell types differed between ROI types within the lesion and at the edge of the lesion (**Fig. 5C, D, G, H**). Notably, the most distinct repositioning of immune cells occurred when tissue-specific ROIs were compared separately (**Fig. 5F-H**). Consistent with the transcriptomics results above (**Fig. 4**), the edge of the lesion showed the most significant changes between the *Mtb*-only and the anti-CD4 mAb treated groups. Most notably, the overall increase in B cells following anti-CD4 mAb treatment previously observed by multiplex IHC was localised to the edge of the lesion (**Fig. 5H**) and was associated with a parallel marked decline of *Cd19* within the lesion (**Fig. 5G**). *Cd8* increased at the edge of the lesion in the BCG-vaccinated group, and *Itgam* (CD11b), and *Itgax* (CD11c) increased in the anti-CD4 mAb reactivation group. Taken together, these transcriptomic results further support a repositioning of non-CD4 immune cells at the edge of the lesion in response to vaccination and depletion of CD4^+^ T cells.

**Figure 5.**
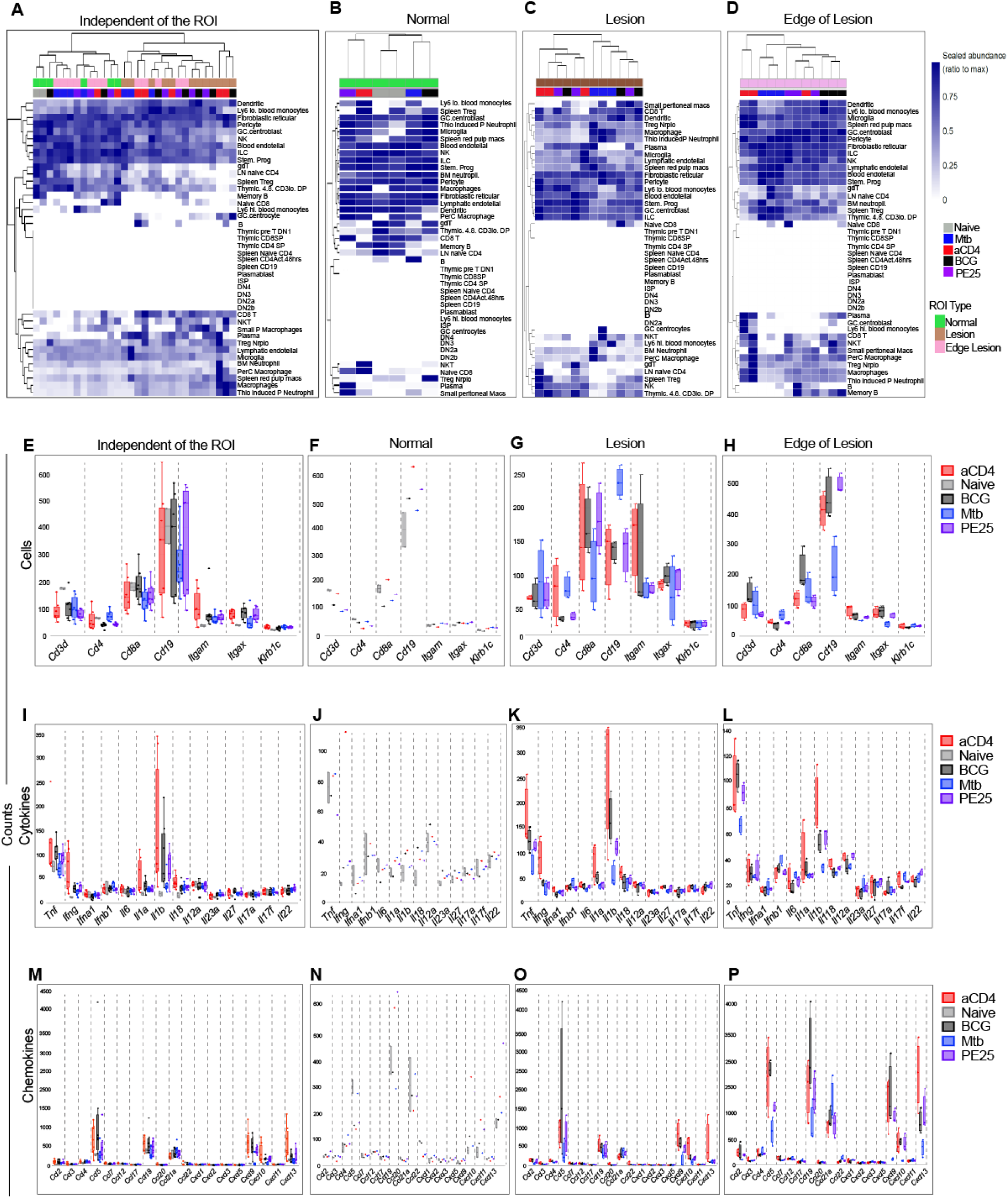
Cellular deconvolution confirms distinct cellular patterns during containment and reactivation of LTBI. **(A-D)** Heatmaps showing the scaled abundance of different cell types identified by cellular deconvolution in all ROIs **(A)** and grouped into tissue locations **(B-D). (E-H)** Gene transcript-based identification of cell subsets with estimated proportions of *CD3*, *CD4*, *CD8a*, *CD19*, *Itgam* (CD11b), *Itgax* (CD11c) and *Klrb1c* (NK) positive cells (aligned with cell populations analysed by Opal Multiplex IHC) across all ROIs **(E)** and depending on the ROI in different tissue locations: normal (**F**), lesion (**G**), and edge of lesion (**H**). **(I-P)** Gene transcript-based identification of key cytokines **(I-L)** and chemokines **(M-P)** involved in *Mtb*-associated immune responses. Results are presented as heatmaps **(A-D)**, box plots ± SEM **(E-P)** from 2 ROIs in normal naïve, 1 ROI in normal in disease from each treatment group, and 3 ROIs in the lesion, and the edge of lesion derived from 7 LNs per experimental group and 2 LNs from naïve controls.

Differences in cytokine (**Fig. 5I-L**) and chemokine (**Fig. 5M-P**) gene expression were also noted across various tissue types when compared to the *Mtb*-only group. In the lesions, the levels of *Ifnγ*, *Tnf*, *Il1b*, *Ccl2*, and *Cxcl19* increased in the anti-CD4 mAb reactivation group. Additionally, *Il1b* also increased in other groups infected with *Mtb*. At the edge of the lesion, *Tnf*, *Il1b*, *Cxcl9*, *Cxcl10*, and *Cxcl13* increased in all groups treated with anti-CD4 mAb. *IL-18*, *Il12a*, and *Ccl2* levels increased, while *Ifn1b*, *Ccl21a*, and *Ccl22* levels decreased in the anti-CD4 mAb reactivation group. In the BCG-vaccinated group, *Ccl5* and *Ccl19* increased, while *Il6* and *Il17f* decreased. Moreover, *IL18*, *Il12a*, *Ccl2*, and *Ccl5* levels increased in the rBCG-vaccinated group, while INF-1β levels decreased. In summary, these findings align with the cellular results showing an altered expression of cytokines and chemokines in the different tissue types, with major differences observed at the edge of the lesion.

### Lethal LTBI reactivation only occurs in the combined absence of lymphocytes and is independent of innate responses

The sum of our analyses thus far inferred that lymphatic LTBI containment may be associated with an expansion and repositioning of immune cells, particularly B cells, CD8^+^ T cells and NK cells at the edge of the LN lesions. To experimentally address the role of these cells for lymphatic LTBI containment, we induced lymphatic *Mtb* infection in a variety of genetically modified mouse strains with and without vaccination. First, we infected wild-type C57BL/6 mice, µMT mice that lack mature B cells [41], *Rag1^-/-^* mice that lack mature B and T cells [42] and *Rag2^-/-^IL2rg^-/-^* mice that lack mature B, T and NK cells [43] i.d. with *Mtb*. Two groups of C57BL/6 mice received anti-CD4 mAb treatment either alone or in addition to *Mtb* infection and naïve mice were also included as controls (**Fig. 6A**). While all C57BL/6 mice and all µMT mice survived *Mtb* infection for 120 days (**Fig. 6 B, C**), 100 % of infected *Rag1^-/-^* mice succumbed to the infection with a median survival of 71 days (**Fig. 6D**). Severely immunocompromised *Rag2^-/-^IL2rg^-/-^* mice only survived for an average of 56 days (**Fig. 6E**). These results indicated that the absence of CD4^+^ T cells or B cells alone does not lead to death, whereas the combined absence of B and T cells does. The results also implied that NK cells may play a minor contribution to LTBI containment in the absence of B and T cells. As expected, anti-CD4 mAb-treated C57BL/6, *Rag1^-/-^* and *Rag2^-/-^IL2rg^-/-^* mice also showed high levels of CFU in the LN, lung and spleen (**Fig. 6F-H**) as well as significantly elevated histopathological damage in LNs (**Fig. 6I, J**) and lungs (**Fig. 6K, L**). However, when we enumerated the *Mtb* burden in surviving C57BL/6 and µMT mice, we were intrigued to observe that µMT mice showed almost no signs of bacterial spread nor a productive infection in the lymphatics (**Fig. 6F-L**). RNA sequencing of fixed LN tissue also revealed that the transcriptomic response of µMT mice to lymphatic *Mtb* infection differed from WT C57BL/6 as well as from immunocompromised *Rag1^-/-^* and *Rag2^-/-^IL2rg^-/-^*mice (**Fig. 6M, N**). Furthermore, pathway and regulator analysis of RNAseq data also revealed that severe infection in immunocompromised *Rag1^-/-^*and *Rag2^-/-^IL2rg^-/-^* mice and anti-CD4 mAb-treated C57BL/6 mice is associated with a particularly strong neutrophil degranulation signature that is not visible in µMT mice. Instead, µMT mice were associated with CTLA4 signalling in cytotoxic T lymphocytes **(Fig. S9A, B)**.

**Figure 6.**
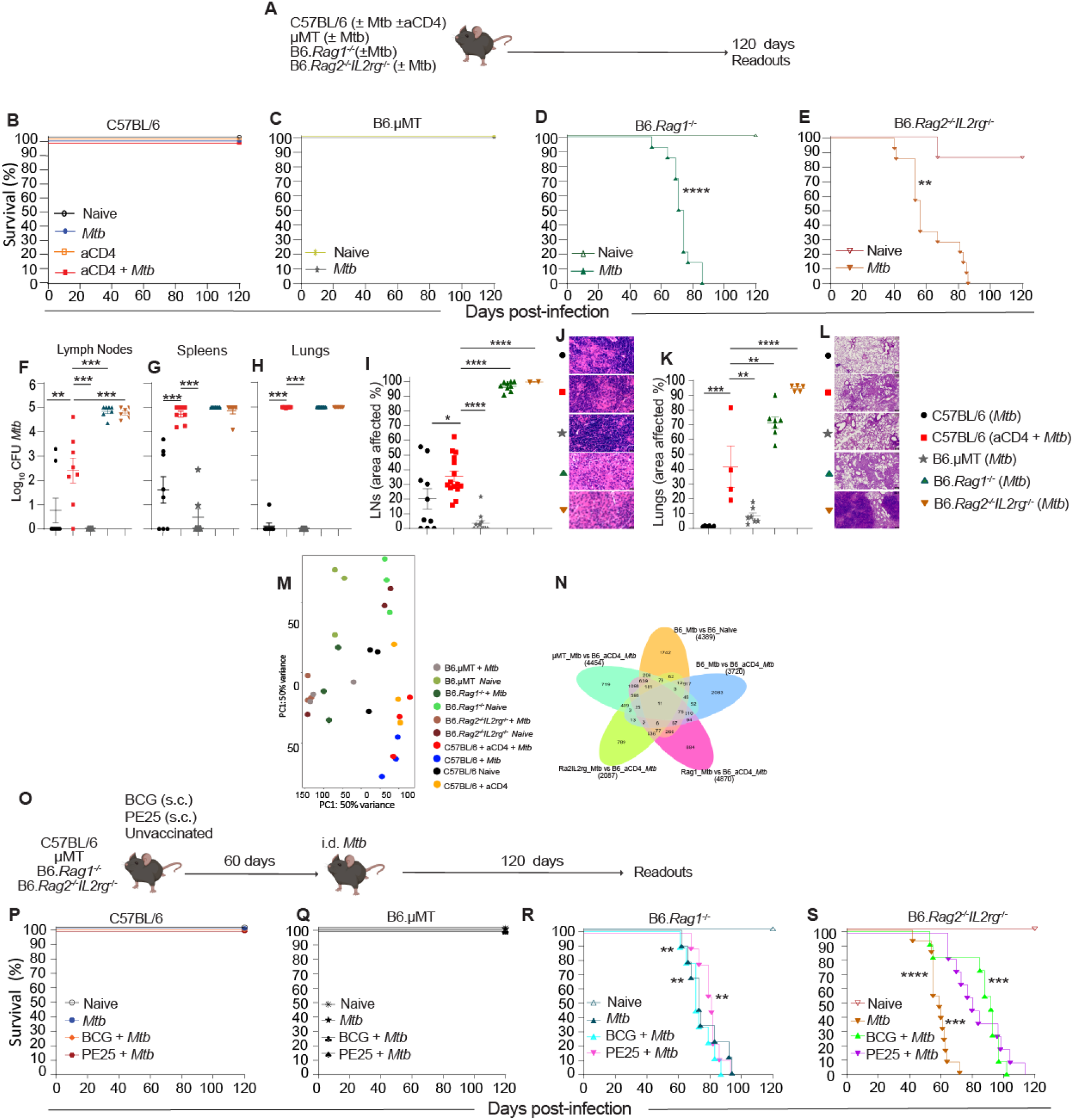
Lethal LTBI reactivation occurs in the combined absence of lymphocytes and is independent of innate responses. **(A)** Experimental setup of unvaccinated mice. (**B-E)** Percent survival of C57BL/6 **(B)**, µMT **(C)**, *Rag1^-/-^* (D) and *Rag2^-/-^IL2rg^-/-^* (E) mice following i.d. infection with *Mtb* and anti-CD4 mAb treatment (C57BL/6 only; **B**). (F-H) *Mtb* CFU burden in cervical LNs **(F)**, spleens **(G),** and left lung lobes **(H)** at the time of euthanasia or at 120 days after *Mtb* infection for surviving mice. Percent of tissue affected by *Mtb* in cervical LNs **(I)** and left lobe lungs **(K)**. Representative H&E sections of cervical LNs **(J)** and left lung lobes **(L)**. **(M-N)** Bulk-RNA sequencing results obtained from paraffin-embedded cervical LN sections. **(M)** PCA map of RNAseq of wild-type, anti-CD4 mAb immunosuppressed, and mutant mice. Venn diagram comparing gene expression of infected mutant mice with C57BL/6 groups **(N)**. **(O)** Experimental setup of mice following i.d. infection with *Mtb* and prior vaccination with BCG or BCG::ESAT-6-PE25SS. (**P-S)** Percent survival of C57BL/6 **(P)**, µMT **(Q)**, *Rag1^-/-^* (R), and *Rag2^-/-^IL2rg^-/-^* (S). Results are presented as individual data points ± SEM **(F-I, K)**, representative H&E images **(J, L)**, and Kaplan-Meier survival curves **(B-E, P-S)**. Data pooled from 2-3 independent experiments. Statistical analyses: One-way ANOVA followed by Tukey’s multiple comparison test **(F-I, K)** and Long-rank (Mantel-Cox) test for survival curves **(B-E, P-S)**. Significant differences are indicated by asterisks: * *p* < 0.05, ** *p* < 0.01, *** *p* < 0.001, **** *p* < 0.0001. See also Figure S9.

We reasoned that if BCG vaccination-mediated containment of LTBI was driven by innate immune mechanisms, then vaccination of *Rag1^-/-^*or *Rag2^-/-^IL2rg^-/-^* mice before *Mtb* infection should extend their survival (**Fig. 6O**). However, vaccination with BCG or BCG::ESAT-6-PE25SS did not extend the survival of *Rag1^-/-^*mice (**Fig. 6R**) and only marginally extended the survival of *Rag2^-/-^IL2rg^-/-^* mice (**Fig. 6S**). Also, vaccination did not change the survival of µMT or C57BL/6 mice (**Fig. 6P, Q**). Collectively, these results indicated that prevention of severe disease and death after LTBI reactivation may either be driven by B and T cell interactions or by a compensatory effect of CD8^+^ T cells. The results also demonstrated that vaccine-mediated LTBI containment is driven by adaptive immune responses with only a minor contribution of innate immunity in the absence of adaptive immune cells.

### CD8^+^ T cells mediate LTBI containment following anti-CD4 mAb treatment and B cells are dispensable

Next, we wanted to further quantify the relative contributions of B cells and different T cell subsets in the containment of LTBI. For this, we first established an adaptive transfer system that would allow us to dissect the contributions of cell subsets on mouse survival (**Fig. 7A**). We confirmed that adaptive transfer of LN-or spleen-derived cells from wild type C57BL/6 mice into *Rag2^-/-^ IL2rg^-/-^* mice can reverse the 100% mortality observed after *Mtb* infection into 100% survival (**Fig. 7B**). This provided a tractable system to transfer LN cells that were depleted of either T cells, B cells or both (**Fig. 7C**). Almost all mice that received total LN cells, regardless of whether cells came from naïve or vaccinated donors, survived for 120 days (**Fig. 7D**). In line with the results seen in µMT mice, the survival data clearly showed that the absence of B cells from transferred cells can be compensated for, with 100% of mice that received cells from B cell depleted vaccinated animals surviving for 120 days and approximately 70% of mice that received cells from B cell depleted naïve mice surviving for 120 days (**Fig. 7E**). In contrast, mice that received either CD3^+^ (**Fig. 7F**) or CD3^+^ and CD19^+^ cell-depleted LN cells (**Fig. 7G**) succumbed to the reactivation of LTBI between 40 and 80 days, which was indistinguishable from *Rag2^-/-^IL2rg^-/-^*mice that had received no cells. These results mirrored the survival data seen in *Rag1^-/-^* mice (**Fig. 6D**) but pointed towards a critical role for T cells rather than B cells in mediating LTBI containment.

**Figure 7.**
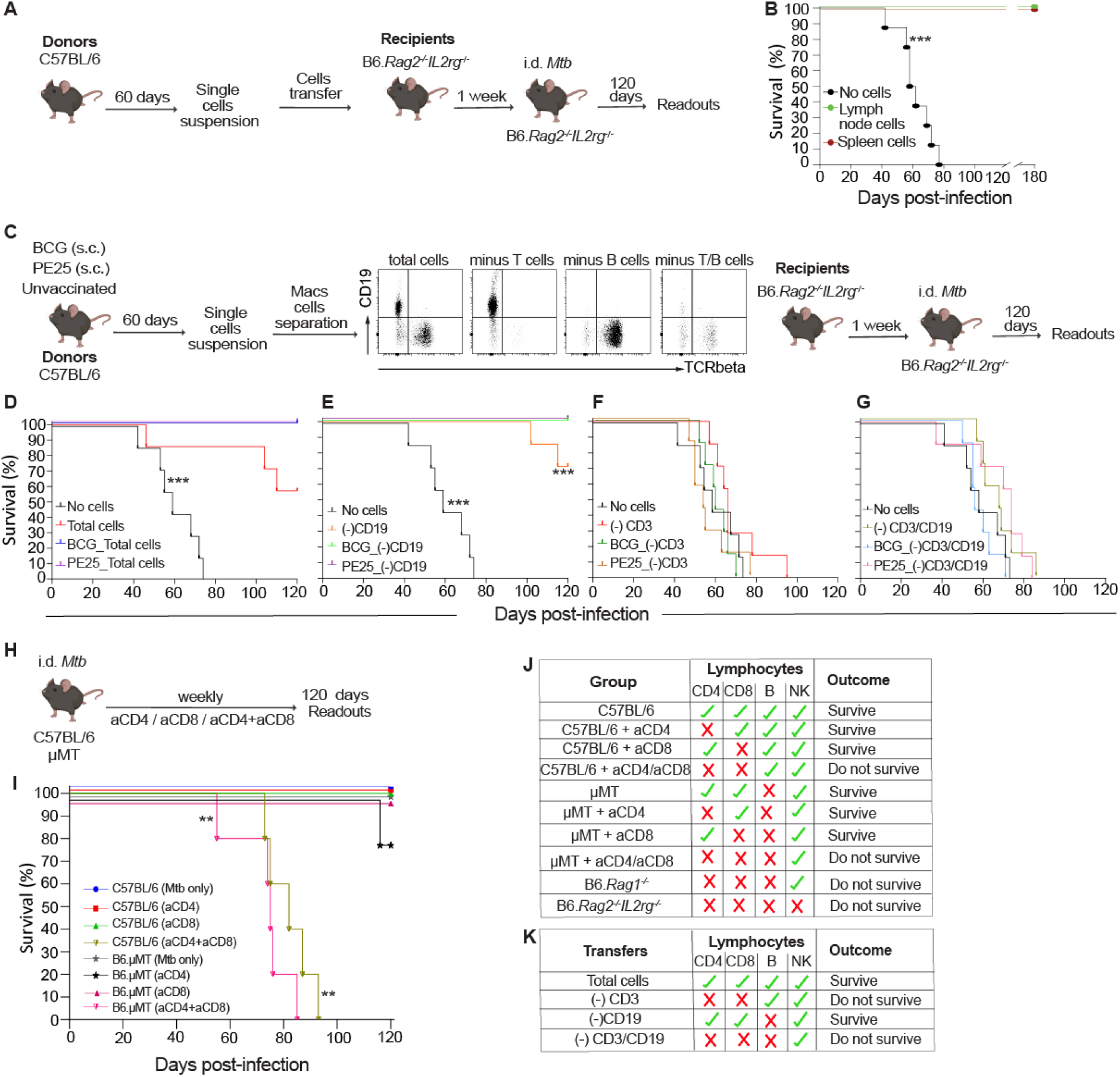
CD8^+^ T cells mediate LTBI containment following immunosuppression and B cells are dispensable. **(A)** Experimental setup of mice receiving adoptive cell transfer (Recipients) from spleen and LNs derived from naïve C57BL/6 mice (Donors). **(B)** Percent survival of *Rag2^-/-^IL2rg^-/-^* mice following i.d. infection with *Mtb* that have received either LN cells, spleen cells, or no cells from naïve donors. **(C)** Experimental setup for adoptive transfer from unvaccinated and vaccinated mice (Donors), including representative FACS plots of LN cells following depletion of T cells, B cells, or both. **(D-G)** Percent survival of *Rag2^-/-^IL2rg^-/-^* mice (Recipients) following i.d. infection with *Mtb* that have received either total LN cells (**D**), B cell-depleted LN cells (**E**), T cell-depleted LN cells (**F**), or B and T cell-depleted LN cells (**G**) from naïve donors or BCG or BCG::ESAT-6-PE25SS-vaccinated donors. **(H)** Experimental setup of C57BL/6 and µMT mice treated weekly with anti-CD4 mAb, anti-CD8 mAb or both anti-CD4/anti-CD8 mAb. **(I)** Survival of C57BL/6 and µMT treated with anti-CD4 mAb, anti-CD8 mAb or both anti-CD4/anti-CD8 mAb. **(J)** Table summarising the survival results of all wild-type and knockout mice with and without immunosuppression via mAb-mediated cell depletion. **(K)** Table summarising the survival results of *Rag2^-/-^IL2rg^-/-^*mice receiving total LN cells, or T cell, B cell or T and B cell-depleted LN cells. Results are presented as survival means (**B, D-G, I**) and representative FACS plots **(C)**. Statistical analyses: Long-rank (Mantel-Cox) test for survival curves. Significant differences are indicated by asterisks: ** *p* < 0.01, *** *p* < 0.001.

To definitively assess the relative contributions of CD4^+^ T cells, CD8^+^ T cells and B cells to the prevention of lethal LTBI reactivation, we performed an additional experiment in which C57BL/6 wild-type mice and B cell-deficient µMT mice were treated with anti-CD4 mAb, anti-CD8 mAb or both (**Fig. 7H**). Strikingly, in both genetic backgrounds only mice treated with both anti-CD4 and anti-CD8 mAbs succumbed to the infection (**Fig. 7I**), with similar kinetics to *Rag1^-/-^* mice (**Fig. 6D**) and *Rag2^-/-^IL2rg^-/-^*mice that had received T cell-depleted LN cells (**Fig. 7E**). Taken together, our results clearly demonstrate that LTBI containment is driven by T cells and that B cells are dispensable for this process (**Fig. 7J, K**). While the absence of either CD4^+^ or CD8^+^ T cells can be compensated for, regardless of whether B cells are present or not (C57BL/6 vs. µMT), the combined absence of both CD4^+^ and CD8^+^ T cells was always lethal. Our results also definitively show that NK cells alone cannot compensate for the loss of T cells.

Collectively, our data provide a comprehensive assessment of the cellular requirements for the containment of otherwise fatal lymphatic LTBI reactivation and provide strong evidence that CD8^+^ T cell-mediated immunity should be targeted to avoid LTBI reactivation when CD4^+^ T cell immunity is compromised, such as in the context of HIV co-infection.

## Discussion

Despite the availability of a licensed vaccine for over 100 years, TB still claims over a million lives per year, and it is widely accepted that a lack of clear immunological correlates of protection has prevented the development of a better TB vaccine [44]. Hence, the identification of such correlates remains the ‘holy grail’ in TB research. Given that a proportion of active TB is a consequence of reactivated LTBI [45], understanding the mechanisms that control *Mtb* containment will likely also contribute to the development of an improved vaccine and potential immunotherapeutic approaches for LTBI.

Here, we provide compelling evidence that contained lymphatic infection of *Mtb* is exclusively maintained by hierarchical and redundant contributions of T cells. While deficiency in either CD4^+^ T cells or CD8^+^ T cells led to a slow but progressive reactivation of LTBI, rapid and lethal reactivation only occurred in the absence of both T cell subsets. In contrast, our results show that CD4^+^ and CD8^+^ T cell deficiency could not be compensated for by B cells and that B cells are not required for lymphatic containment in this model of LTBI. These findings provide detailed insights into the reactivation dynamics of LTBI and provide concrete evidence that CD8^+^ T cells are sufficient to control lymphatic *Mtb* infection. Our results also show that BCG/rBCG vaccination-mediated reduction, and in some cases even prevention of LTBI reactivation, is a consequence of the repositioning of CD8^+^ T cells and B cells at the edge of lymphatic lesions. While B cells appear to be passive bystanders, CD8^+^ T cells actively promote LTBI containment in the infected LNs rather than via preferential killing in the lung.

It has previously been shown that in certain situations, B cells can contribute to the immune response against *Mtb* via antibody and cytokine production as antigen-presenting cells and through their contributions to the formation of granulomas [46]. Surprisingly, the absence of mature B cells in µMT did not lead to LTBI reactivation nor the premature death of infected mice. In fact, µMT mice showed almost no signs of a productive infection with *Mtb* at all. While these results support previous studies that reported that B cells are either not required to control TB or promote detrimental effects [47, 48], they also contradict others in which B cells and antibodies played an important role in TB immunity [49–51]. It is increasingly clear that the role of B cells in TB is context-and animal-model dependent [46, 52]. The findings obtained in our study imply that B cells are either actively attracted and exploited by *Mtb* to establish an early, productive infection in the LN, or that B cells are predominantly needed to help T cell localisation within the granuloma. This hypothesis is supported by results from an elegant study showing that *Mtb* infection leads to a massive influx of B cells into the lung draining mediastinal LNs, which limits protective T cell activation [53]. It is also conceivable that the expansion of B cells seen in the LNs after CD4^+^ T cell depletion is simply a consequence of the increase in antigen and cell-free *Mtb* bacilli after the breakdown of the granuloma [54].

Upon reactivation of lymphatic LTBI with anti-CD4 mAb we observed a strong neutrophil degranulation signature. Given that neutrophilia is a common response to bacterial infections [55], and neutrophils are generally not part of the long-term structure of the TB granuloma, the degranulation signature likely reflects the granuloma breakdown and increased bacterial burden associated with the anti-CD4 mAb-mediated granuloma disruption [56]. Very recently, it has also been shown that antibodies with distinct Fc variants could restrict *Mtb* growth via a neutrophil-dependent pathway [57]. It is also known that B cells regulate neutrophilia during *Mtb* infection and BCG vaccination [58]. Given that our spatial transcriptomics results highlighted both neutrophil degranulation as well as antibody responses as the top hits associated with LTBI reactivation, it is possible that B cell-neutrophil interactions can also further amplify *Mtb* infection in the lymphatics. It is also conceivable that the increase in lymphatic B cells and neutrophils serves as a host compensatory mechanism to overcome CD4^+^ T cell depletion. Alternatively, B cell expansion could also help to direct T follicular-like helper cells into lymphoid follicles, as has recently been shown following aerosol *Mtb* infection [59].

The use of genetic deficiency of T, B or NK cells, combined with adoptive transfer experiments into lymphocyte deficient *Rag2^-/-^Il2rg^-/-^*mice, unequivocally showed that vaccine-mediated containment of *Mtb* in the lymphatics following depletion of CD4^+^ T cells is driven by adaptive immunity, with a very minor (if any) contribution of innate immune mechanisms. In particular, our data support a model whereby CD4^+^ T cell deficiency is compensated for by CD8^+^ T cells. It has previously been shown that the reactivation of latent pulmonary TB induced by antibiotic therapy in mice is exacerbated when CD8^+^ T cells are depleted [60]. It is also known that CD8^+^ T cells can maintain the balance between *Mtb* control and the prevention of excessive tissue damage during LTBI [61].

It is increasingly clear that CD8^+^ T cell immunity should be induced by a successful TB vaccine. Our results show that BCG vaccination leads to a repositioning of CD8^+^ T cells around *Mtb* bacilli in infected LNs, which prevents bacterial escape from the lesion. Given that the expansion of CD8^+^ T cell numbers was only minor, it is likely that CD8^+^ T cell-dependent bacterial control is a consequence of altered functionality. These results provide further support for the development of vaccines that directly target CD8^+^ T cell-driven immune responses. Very recently, it has been shown that CD4^+^ T cells and CD8α^+^ lymphocytes, but not CD8αβ T cells, are crucial for the near-sterile protection offered by intravenous BCG vaccination in non-human primates [62]. The results pointed towards a unique, non-redundant role for CD4^+^ and CD8α^+^ lymphocytes, with depletion of either cell type leading to increased *Mtb* bacterial load in the lungs. Given that we used a CD8α targeting anti-CD8 mAb clone (2.43), and CD8α depletion affects both adaptive and innate-like immune cells, including NK cells and MAIT cells, it is possible that innate-like CD8^+^ cell subsets may also play a role in our model. Although *Rag*-deficient mice lack αβ-positive T cells [42], and *Rag*-deficient mice phenocopied the lethal LTBI reactivation seen with anti-CD8/anti-CD4 mAb treatment, further studies should investigate the precise contribution of CD8α^+^ vs CD8αβ^+^ cells in reactivation of LTBI, and also how these cells mediate their effector functions.

Our study also provides detailed insights into the reactivation dynamics of lymphatic LTBI across a spectrum of organ systems. It appears that anti-CD4 mAb-mediated reactivation of LTBI affects not only multiple peripheral lymphatic sites, such as the cervical and mesenteric LNs and the spleen but also the liver and the bone marrow. Interestingly, BCG vaccination prevented histopathological changes associated with LTBI reactivation in all those reactivation sites. Given that three distinct recombinant strains of BCG prevented LTBI reactivation, it appears likely that the underlying mechanisms may be more broadly applicable to all live attenuated TB vaccine candidates. However, how translatable are those findings to human LTBI? Epidemiological data suggest that BCG vaccination has limited efficacy in preventing human LTBI reactivation [63, 64]. It is, however, important to note that LTBI reactivation in humans predominantly happens in adults, especially in the elderly and after immunosuppression, often many decades after BCG vaccination. As such, it will be important to understand if the immune mechanisms that mediate prevention of LTBI reactivation in this murine model are long-lived, and if BCG revaccination in adults with LTBI could prevent LTBI reactivation in older age. Our findings also provide hope that vaccination-induced CD8^+^ T cell-mediated immune responses can be incorporated into TB vaccines for use in HIV co-infected individuals.

Our study also has limitations. Firstly, while this model of contained *Mtb* infection is now widely accepted as a useful model to mimic LTBI [14, 15, 65], it uses an i.d. inoculation, a route that is rarely the cause of human LTBI. It will be useful to investigate if CD8^+^ T cells also mediate containment of LTBI in other animal models and in humans. As mentioned above, it will also be critical to dissect if the protective effect is attributable to CD8αα-expressing T cell subsets, as well as innate-like cells. For that reason, depletion studies utilising different mAb clones should be conducted. Furthermore, it remains to be investigated whether CD8^+^ cells mediate the containment of LTBI via antibody-dependent cellular cytotoxicity, specific effector molecules or other mechanisms. Our findings also show that there was no outstanding improvement in the containment of LTBI when rBCG strains were used. Given that most rBCGs only differ from BCG in a very small number of antigens, our study implies that the mechanisms of vaccination-mediated containment are largely driven by factors that are shared between BCG and rBCGs. Future studies should investigate which components of BCG drive containment and if containment can also be induced by subunit TB vaccine candidates.

Finally, while we have attempted to utilise multiple high-resolution histopathological and imaging approaches, we did not perform spatial transcriptomics analyses on a single-cell level. Single-cell spatial technology only just emerged when this study was underway, and future studies should endeavour to use single cell transcriptomic resolution for integration with histological techniques for maximum resolution [66, 67].

In conclusion, the results reported here provide direct evidence of CD8^+^ T cell-mediated control of LTBI and question the role of B cells in lymphatic *Mtb* containment. They also provide a detailed atlas and a transcriptomic resource for lymphatic *Mtb* infection with and without prior vaccination. Our findings have profound implications for the understanding of immunity to TB more broadly and the management of LTBI. The robust model of controlled LTBI reactivation used here may also be relevant to gain further detailed insights into the cellular dynamics after LTBI reactivation across a spectrum of different organs.

## Data Availability

Requests for further information and resources should be directed to and will be fulfilled by the lead contact, Andreas Kupz. This study did not generate new, unique reagents. All data reported in this paper will be shared by the lead contact upon request.

## Code availability

This paper does not report original codes. Any additional information required to reanalyze the data reported in this paper is available in the main text and supplemental information or from the lead contact upon request.

## Acknowledgements

We thank Serrin Rowarth, Kylie Robertson, Bradley Taylor, and Olivia Johnson for their support in the Animal Facility. We thank Erin Roberts, and Roselfina Charol for their assistance in the Histopathology Department, and Leanne Taylor for support with BSL3 work.

This study was funded by NIH grant R01AI161822 to S.S and A.K. The laboratory of A.K. is supported by an NHMRC Investigator Grant (APP2008715). M.A.F is supported by NHMRC Investigator Grant (APP5121190). The funders were not involved in the study design, data collection and the decision to submit the article for publication.

## Author contributions

Conceptualisation: SMH, SS, AK. Methodology: SMH, MK, AH, EG, XT, JT, MM, ZCC, LDP, MD, MR, MAFV, AB, QN, SS, MAF, AK. Investigation: SMH, MK, AH, EG, XT, JT, MM, ZCC, LDP, YP, ET, MD, MR, MAFV, AB, QN, SS, MAF, AK. Manuscript writing: SMH, AK. Manuscript editing: SMH, MK, AH, EG, XT, JT, MM, ZCC, LDP, YP, ET, MD, MR, MAFV, AB, QN, SS, MAF, AK. Funding acquisition: SS, MAF, AK.

## Supplementary Information

**Supplementary Figure 1:**
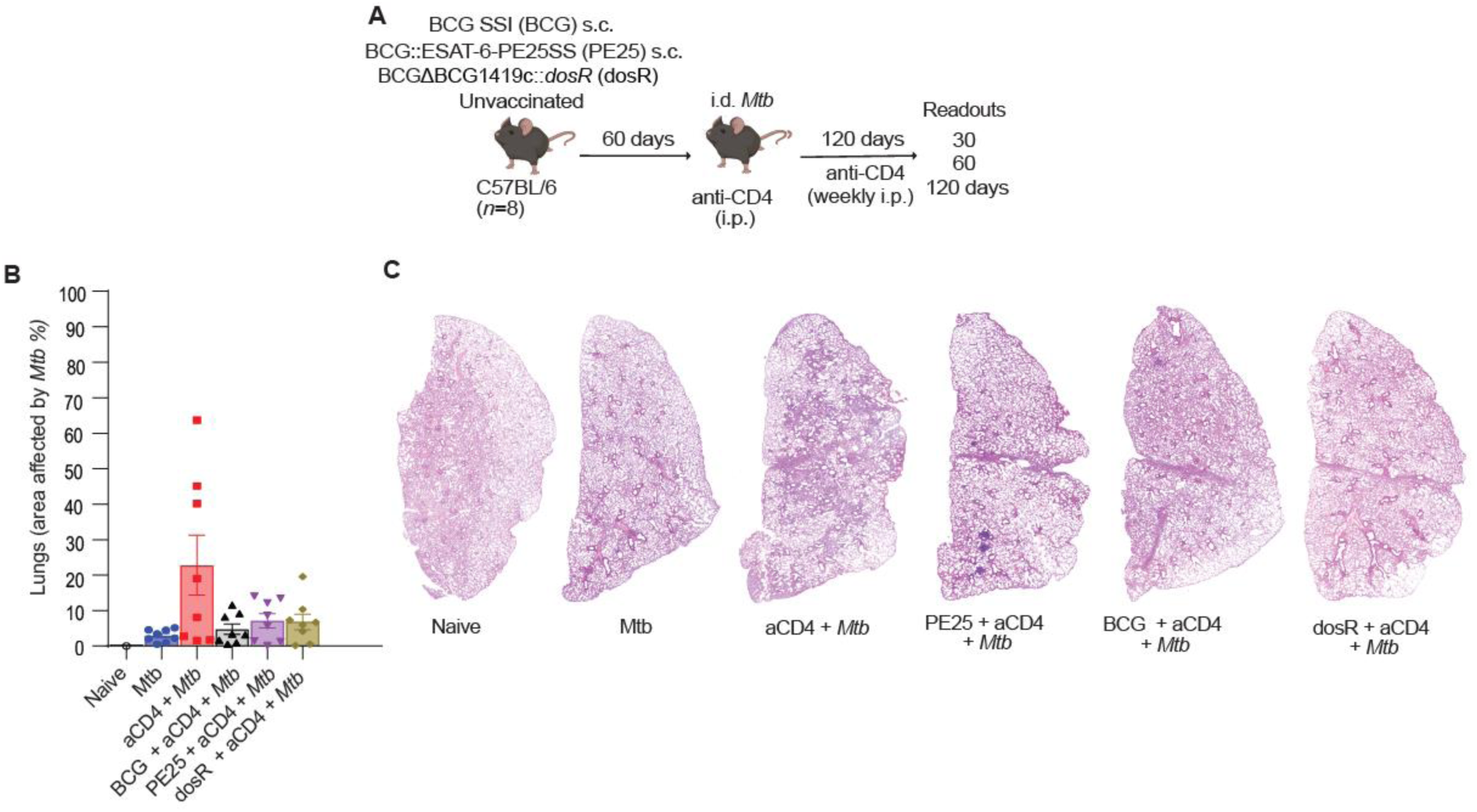
BCGΔBCG1419c::*dosR* vaccination reduces *Mtb*-driven lung pathology following the reactivation of LTBI via immunosuppression. **(A)** Experimental setup. C57BL/6 mice were vaccinated s.c. with 1×10^6^ CFU of either BCG SSI, BCG::ESAT-6-PE25SS, BCGΔBCG1419c::*dosR* or left unvaccinated. Sixty days after vaccination, mice were infected i.d. with 1×10^3^ CFU of *Mtb* H37Rv and received anti-CD4 mAb weekly i.p. for 120 days. **(B)** Percent of tissue affected by *Mtb* in lungs. **(C)** Representative H&E sections of lungs from each treatment group. Results are presented as individual data points ± SEM (n = 8 mice per group).

**Supplementary Figure 2:**
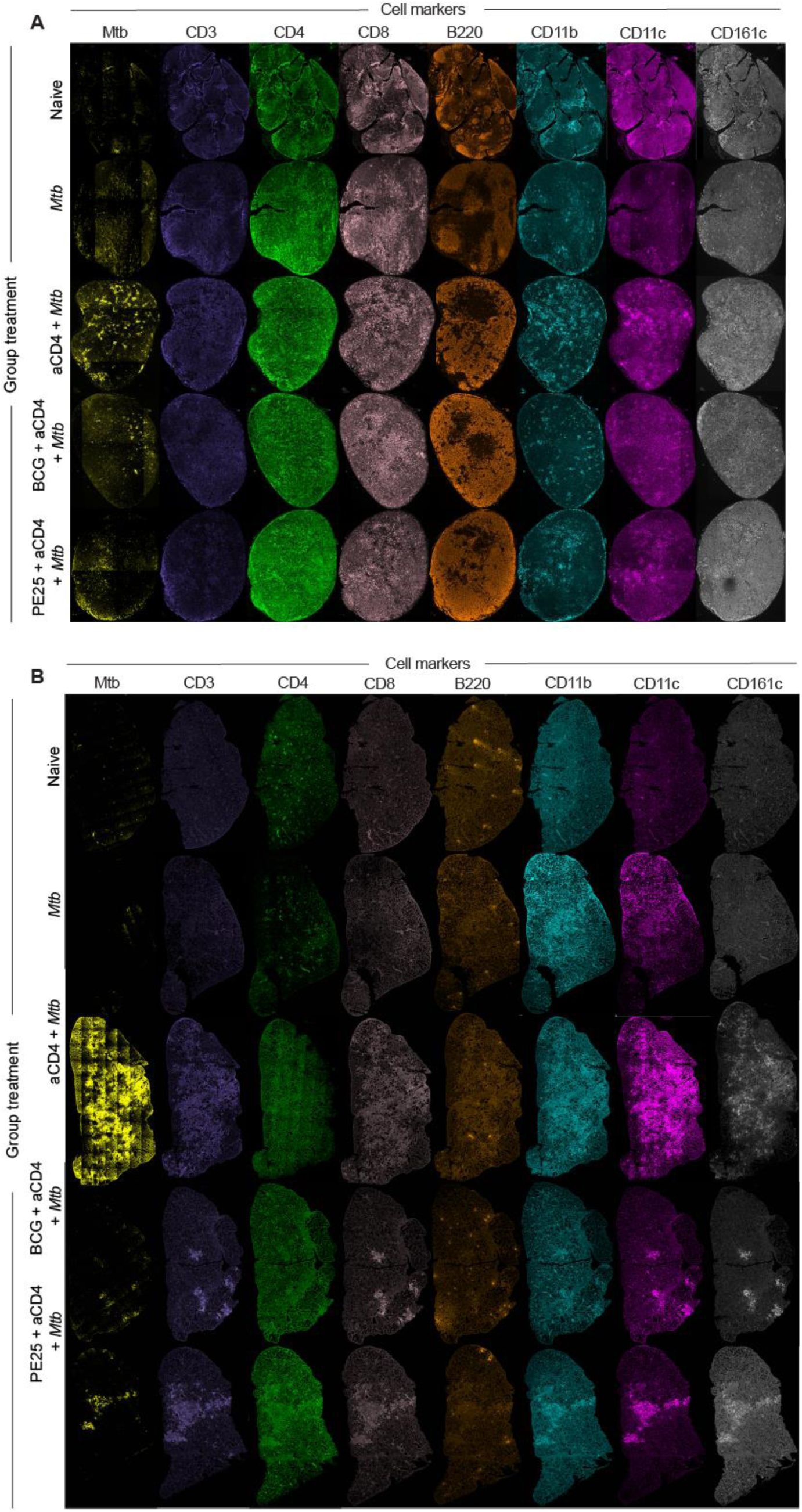
Whole LN and lung sections stained with eight individual fluorescent markers. **(A-B)** Representative images of Opal Multiplex IHC from each treatment group positive for either *Mtb*, CD3, CD4, CD8, B220, CD11b, CD11c or CD161c across entire cervical LNs **(A)** and left lung lobes **(B)**. Results show increased bacterial burden in anti-CD4 mAb treated group and repositioning of immune cells based on vaccination and treatment.

**Supplementary Figure 3:**
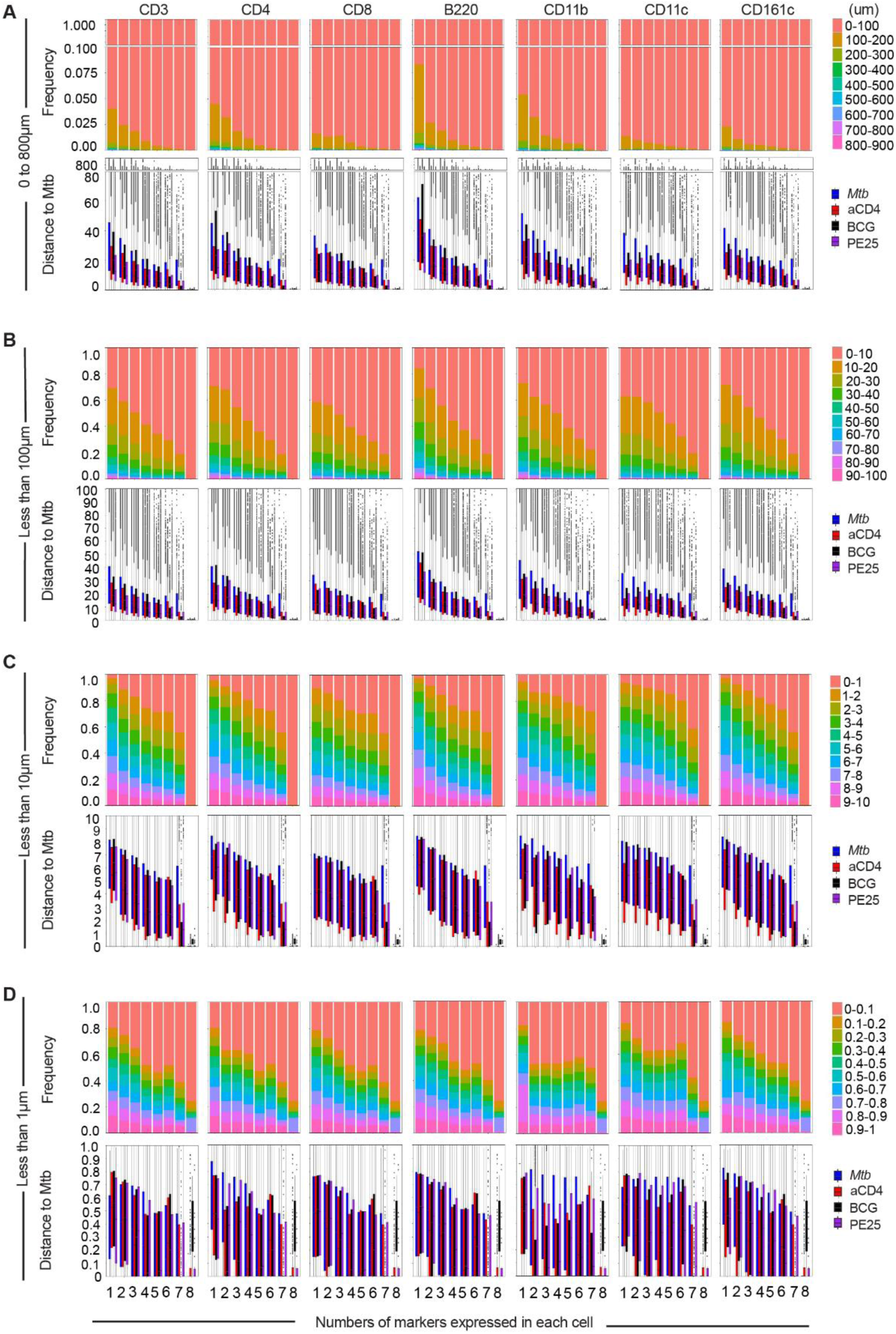
Cell subsets with more than one phenotypic surface marker interact directly with *Mtb*-infected cells but are less frequent. **(A-D)** Frequency of cells in LNs expressing one to eight surface markers and their distance to *Mtb*-infected cells. All cells identified were located at 0-800µm from *Mtb*-infected cells **(A)**, cells situated within 0 to 100µm from *Mtb*-infected cells **(B)**, cells located within 0-10µm from *Mtb*-infected cells **(C)** and cells located within 0-1µm from *Mtb*-infected cells (D).

**Supplementary Figure 4:**
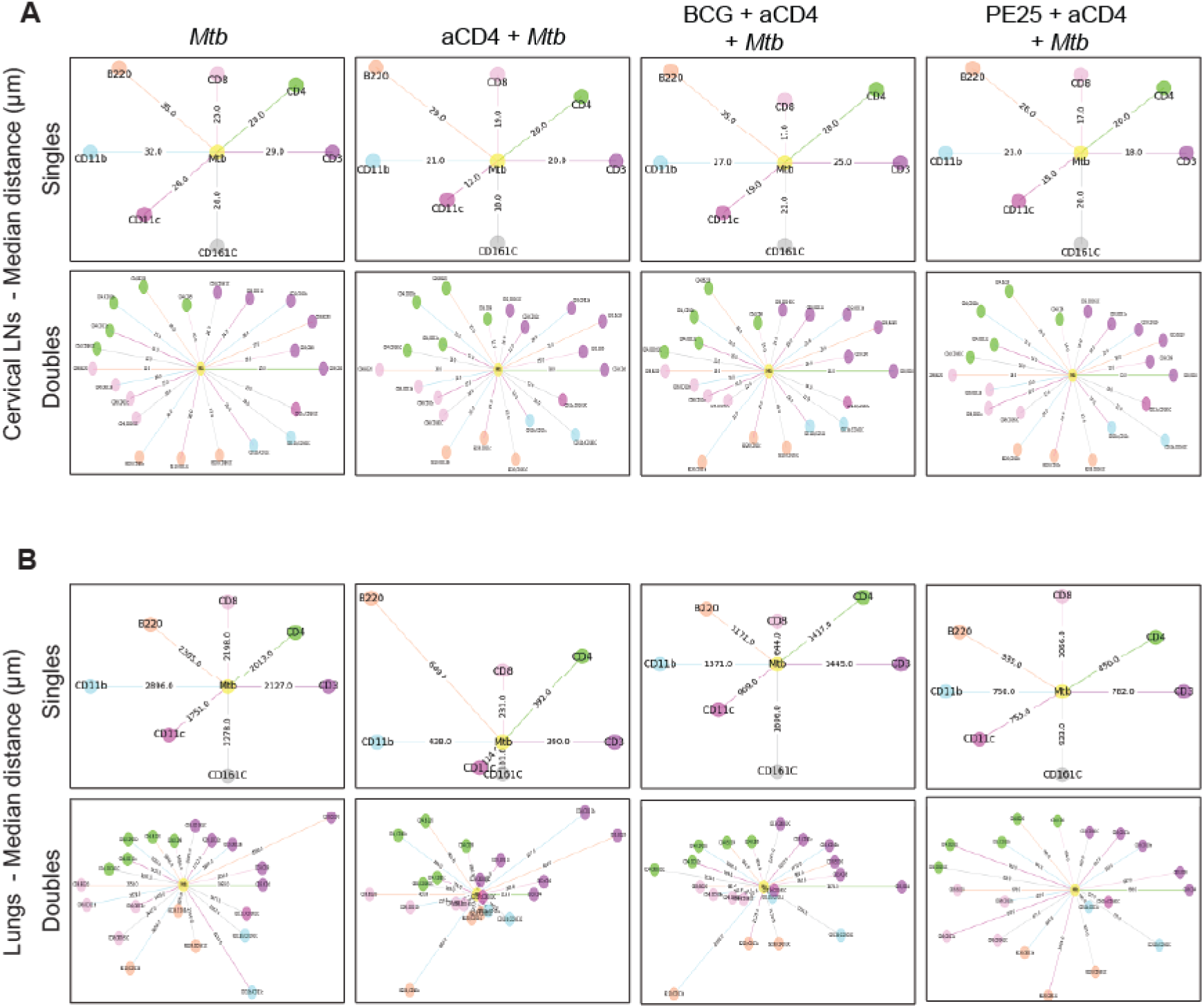
Cell subsets with multiple phenotype markers are in closer proximity to *Mtb*-infected cells. **(A-B)** Median distance to *Mtb*-infected cells in micrometres for cells expressing single or double phenotypic markers in LNs **(A)** and left lung lobes **(B)**.

**Supplementary Figure 5:**
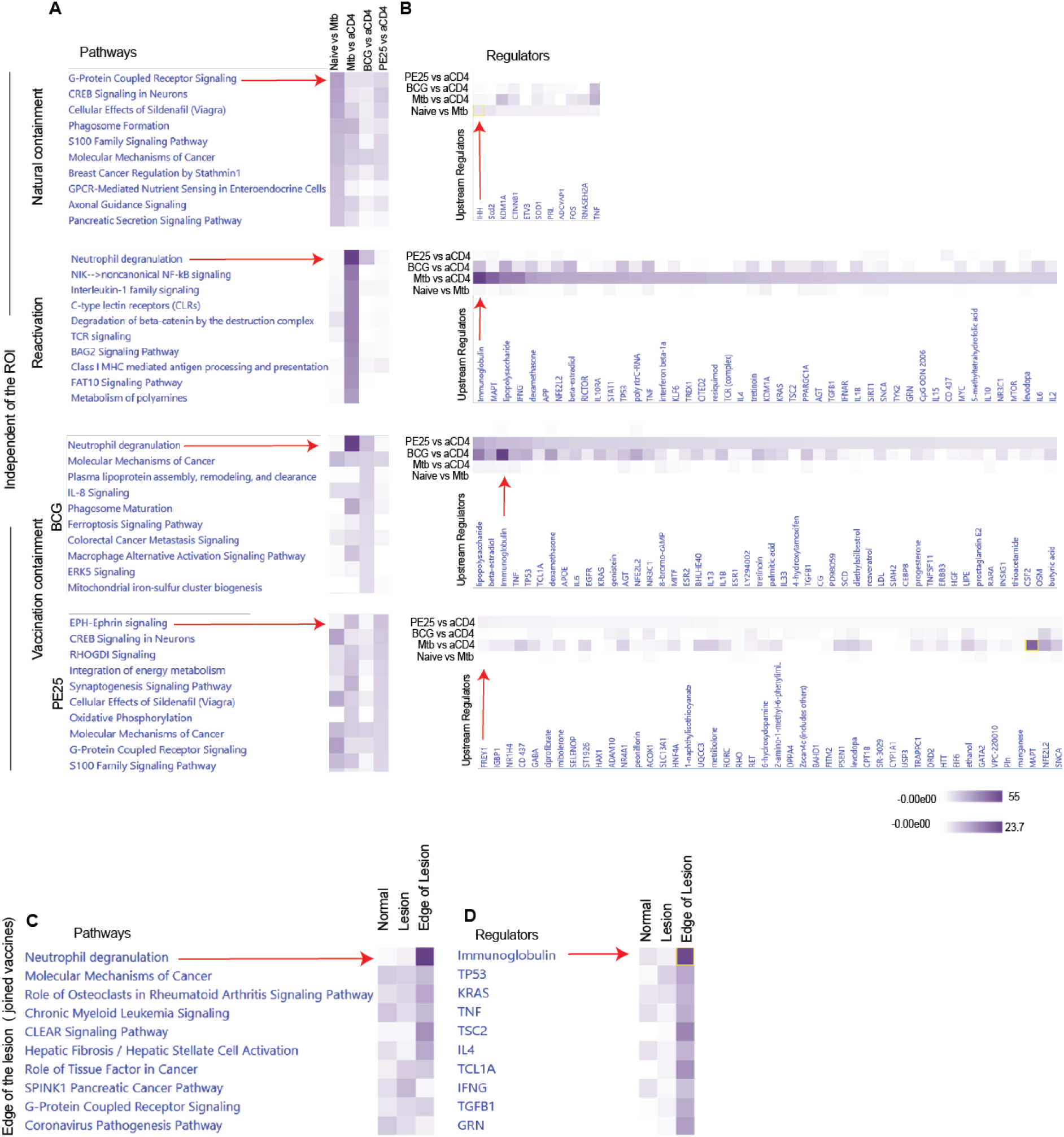
IPA pathways of GeoMx spatial transcriptomics analysis. (**A-D**) IPA pathway and regulators analysis across groups considering the *p* values are consistent with the predicted IPA pathways shown in Figure 4. Pathways **(A)** and regulators **(B)** independent of the ROIs. (**C-D**) Transcriptomic pathways at the edge of the lesion. IPA pathway **(C)** and their regulators **(D)** for combined vaccine-mediated containment groups.

**Supplementary Figure 6:**
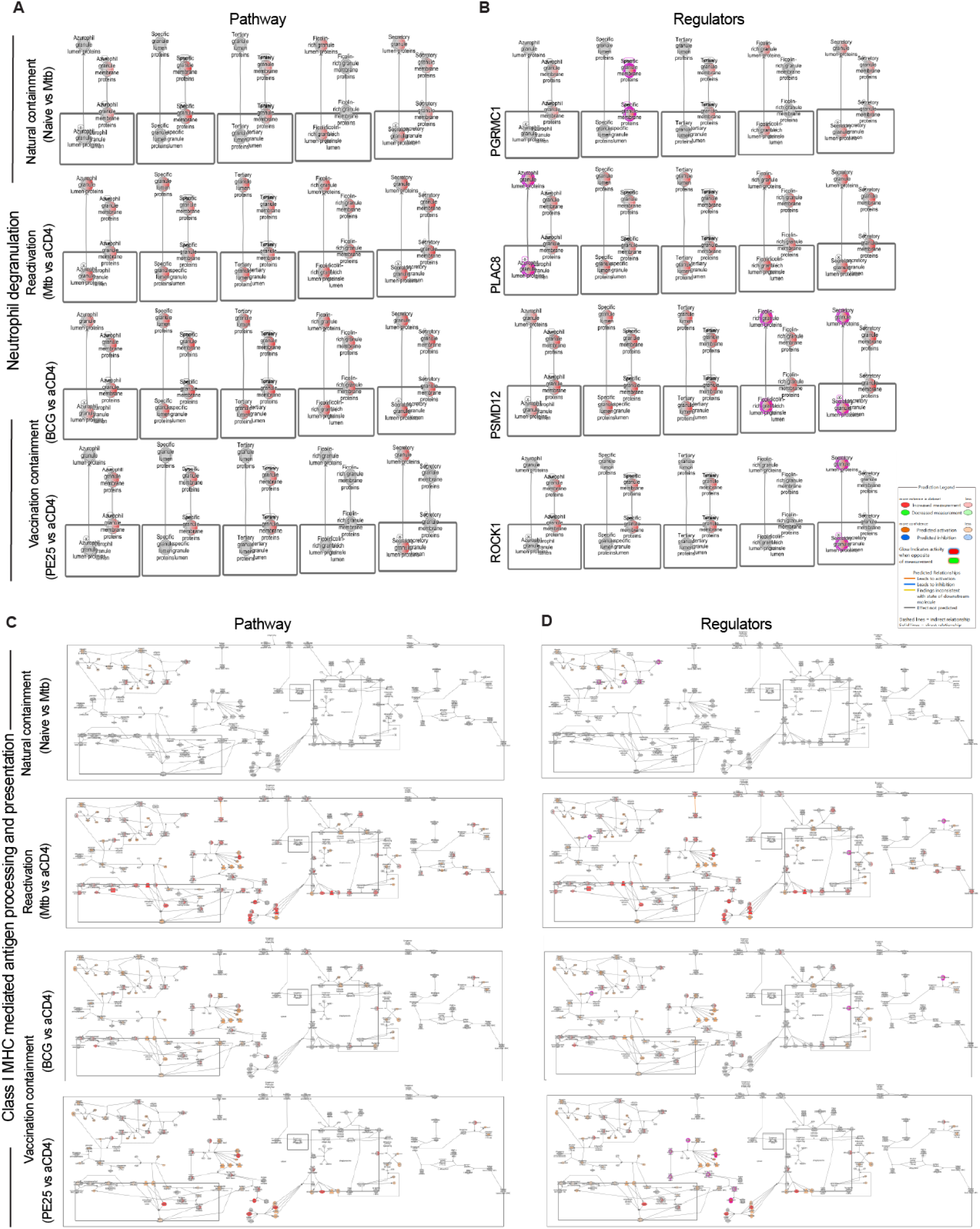
Natural containment, reactivation, and vaccine-mediated containment of LTBI depend on distinct pathways and regulatory circuits. **(A-B)** Gene activity pathways and regulators during neutrophil degranulation. (Left) top pathways characterising different group comparisons: naïve vs. *Mtb*-only; *Mtb*-only vs. anti-CD4 mAb; anti-CD4 mAb vs. each of the vaccines. (Right) expression levels of top-regulated genes across upregulated pathways. **(C-D)** Gene activity pathways and regulators during MHC Class I mediated antigen processing and presentation across natural containment, reactivation, and vaccine-mediated containment. (Left) top pathways characterising different group comparisons. (Right) expression levels of top-regulated genes across upregulated pathways.

**Supplementary Figure 7:**
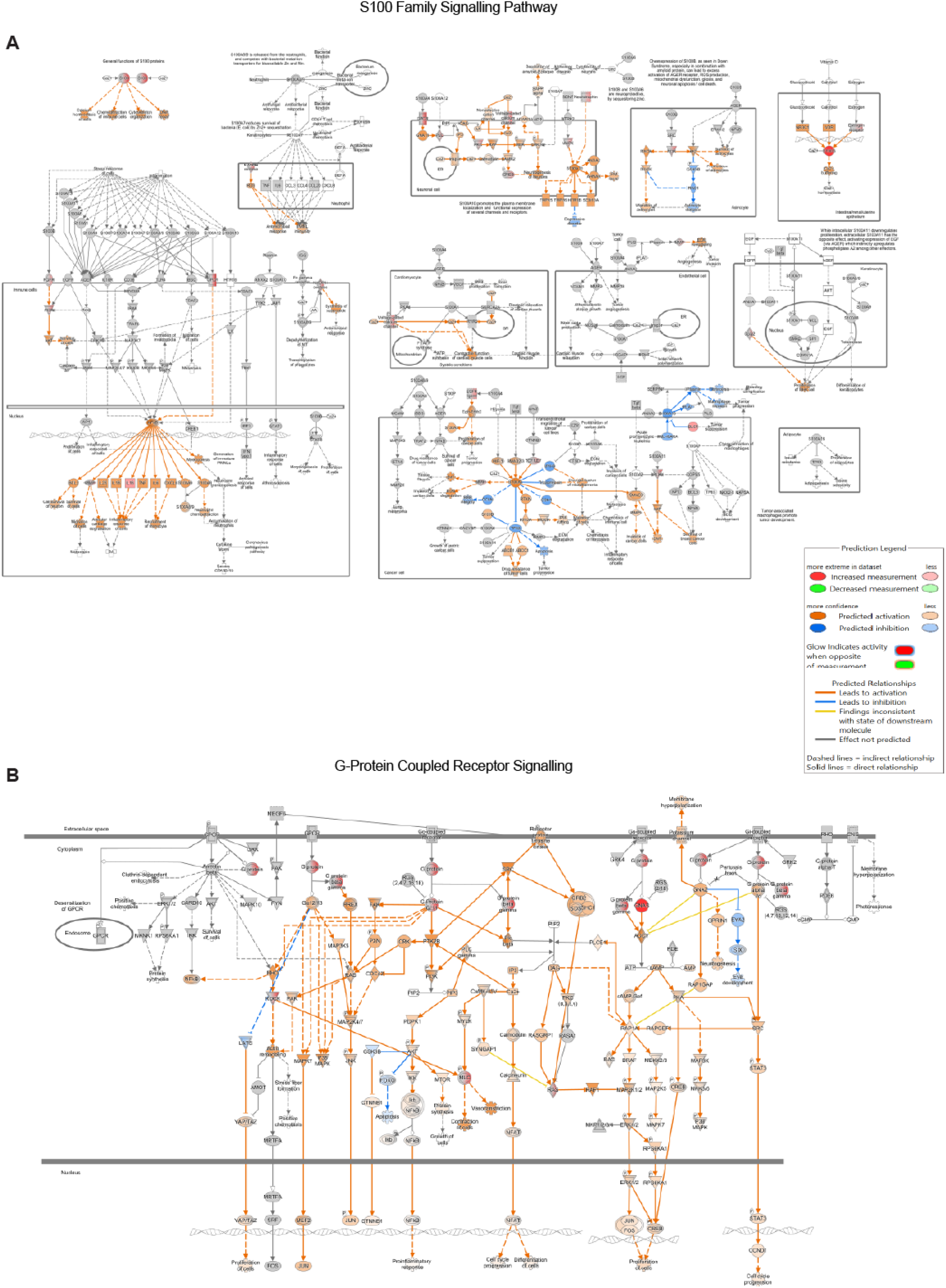
Natural containment of LTBI and protection against LTBI reactivation with BCG::ESAT6-PE25SS may operate through different mechanisms. **(A)** Top pathway characterising natural containment (naïve vs. *Mtb*-only). Pathway identified as S100 family signalling. **(B)** Top pathway characterising BCG::ESAT6-PE25SS-based LTBI containment (*Mtb* + anti-CD4 mAb vs. BCG::ESAT6-PE25SS + *Mtb* + anti-CD4 mAb) Pathway identified as G-protein Coupled Receptor Signalling. Network plots showing predicted up-and down-regulated interactions.

**Supplementary Figure 8:**
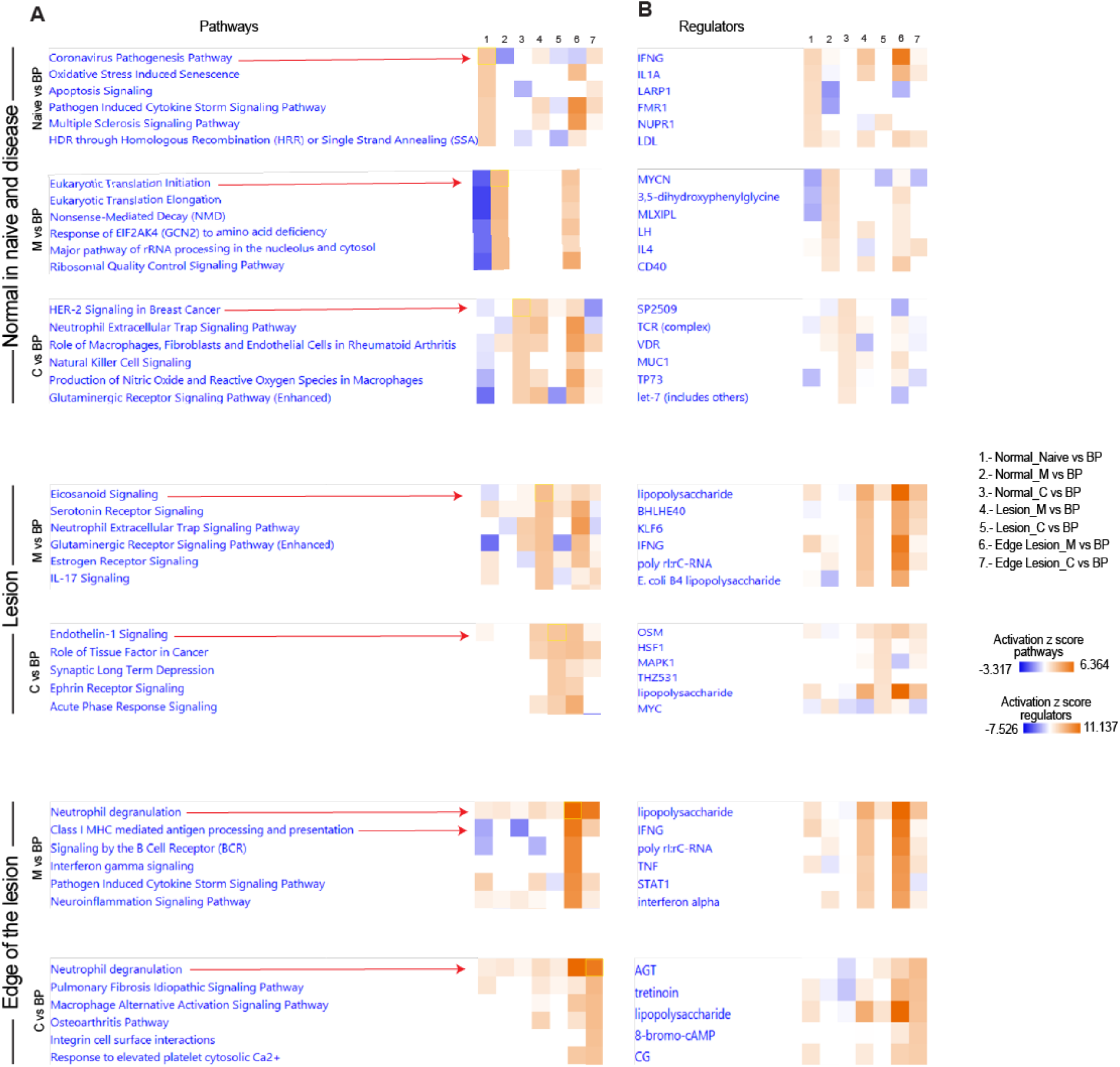
The reactivation, containment, and protection of LTBI at specific ROIs are characterised by unique and shared pathways and regulators. Full data set for Figure 4 P & Q showing spatial transcriptomics-derived IPA predicted pathways **(A)** and their regulators **(B)** at different ROI types across all treatment groups.

**Supplementary Figure 9:**
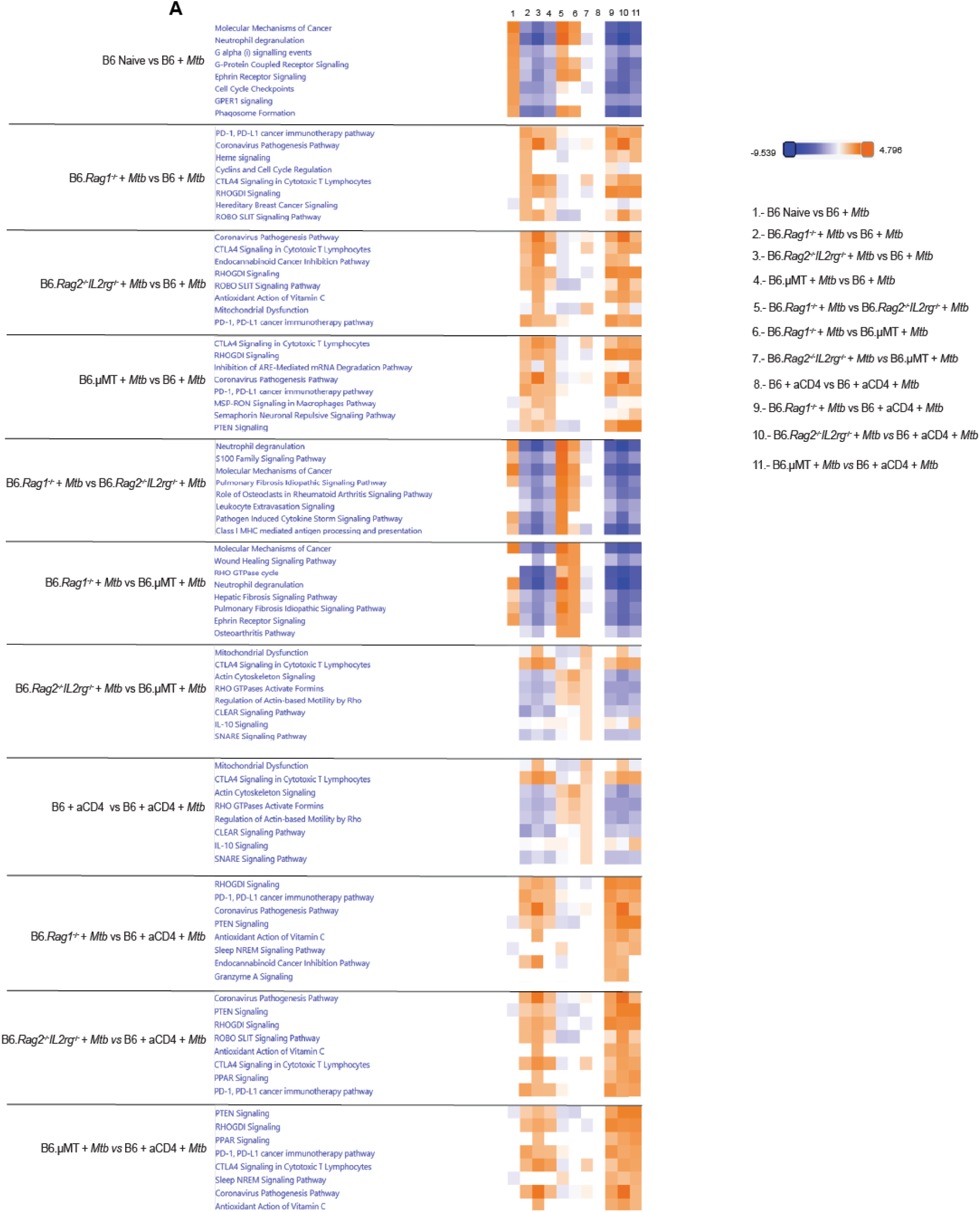
RNAseq-derived pathways and regulators comparing different treatment groups across mouse strains. Bulk-RNA sequencing results obtained from paraffin-embedded cervical LN sections. IPA predicted pathway analysis across mouse strains and treatment groups **(A)**. Results suggest increased activity of cytotoxic T cells in the absence of CD4^+^ T cells and B cells and increased neutrophil degranulation in situations of high bacterial burden as shown in Figure 6F-L.

